# “mRNA deep sequencing profiling and integrative miRNA-mRNA network analysis in pediatric Burkitt leukemia and lymphoma “

**DOI:** 10.64898/2026.10.23.760471

**Authors:** Teresa M. Cardesa-Salzmann, Veronica Celis, Efstathios-Iason Vlachavas, Björn Schneider, Caterina Redwanz, Olia Shokraie, Elena Esperanza-Cebollada, Sara Montesdeoca, Mireia Camós, Silvia Planas, Nuria Conde-Cuevas, Claudia Maletzki, Cristina Jou, Jaume Mora, Almut Meyer-Bahlburg, Carl Friedrich Classen

## Abstract

**Background:** Although the genomic alterations in Burkitt leukemia and lymphoma (BL) have been extensively studied, data on genome-wide dysregulation of messenger RNA (mRNA) and microRNA (miRNA) in Burkitt leukemia (BuAL) and Burkitt lymphoma (BuLY) are scarce. The aim of this study was to analyze mRNA and miRNA expression profiles in pediatric BuAL/BuLY to identify dysregulated mRNA-miRNA networks.

**Methods:** Total RNA from fresh tumor samples of 13 pediatric BL patients (BuAL n=7, BuLY n=6) and 4 pediatric normal control lymphoid tissue (NCLT) samples was included for mRNA-deep sequencing and miRNA-microarrays studies. Differential expression (DE) analysis of mRNAs and miRNAs as well as gene set and functional enrichment analysis were performed. The multiMiR R package was used to identify the validated target genes for the dysregulated miRNAs. miRNA-mRNA target interaction analysis was performed combining DE, validated targets observed among the inverse-significant miRNA-mRNA correlations, with evidence statistics retained (ρ≤−0.5 and Benjamini-Hochberg (BH)-adjusted correlation *p*-value< 0.05).

**Results:** BL showed 1965 DE genes (DEG) compared to NCLT (1214 upregulated, 751 downregulated) (*p*-adj<0.05, |log₂FC|>0.5) and 11 DE miRNAs (three upregulated and eight downregulated) (FC>1.5, p-adj<0.05). The transcriptional differences observed between BL and normal lymphoid tissue inferred by functional enrichment analysis identified along a MYC-driven proliferative backbone with strong E2F4 and MYB pathway activity, activation of the MAPK, PI3K and the G2/M checkpoint signaling pathways and repression of the JAK–STAT, NFkB, TP53, TRAIL and hypoxia pathways (*p*-adj<0.05). Functional enrichment analysis highlighted an upregulation of proliferative and metabolic pathways accompanied by an attenuated inflammatory and immunosurveillance signaling in BL compared to NCLT. Correlated miRNA-mRNA expression analysis identified DE dysregulated miRNAs and their validated target genes. DE downregulated miRNAs (hsa-miR-150-5p, hsa-miR-342-3p and hsa-miR-450-5p, among others) inversely correlated with upregulated mRNA targets showing pathway enrichment related to transcriptional activity and nuclear organization modules. mRNA targets of DE upregulated miRNAs (hsa-miR-106a-5p, hsa-miR-17-5p and hsa-miR-7-5p) were enriched in downregulated pathways related to immune response processes.

**Conclusions:** Overall, this pilot study provides new insights on key proliferative, metabolic and cytokine-mediated immune signaling pathways in BL and highlights miRNA-mRNA networks influencing transcriptional activity and immune surveillance, paving the way for experimental validation models.

## Background

Burkitt lymphoma/leukemia (BL), the most common non-Hodgkin lymphoma subtype in children(1), is a highly aggressive mature B-cell neoplasm with a germinal center B (GCB) phenotype(2, 3) characterized by dysregulation of *MYC* due to the hallmark translocation that places *MYC* expression under the control of one of the immunoglobulin gene loci (4–6).

Burkitt lymphoma (BuLY) and its leukemic counterpart, Burkitt leukemia (BuAL), defined by > 25% bone marrow infiltration, have overlapping clinical and biological features and are considered different presentations of a single disease. While treatment with current risk-adapted chemo-immunotherapy achieves an event-free survival of 90-95% in pediatric cohorts(7, 8), relapsed and refractory BL have a dismal prognosis, with survival rates below 30%(9). Therefore, further elucidating mechanisms involved in BL pathogenesis is of upmost importance. Despite the presence of *MYC* translocations in all BL, cooperating genomic alterations are required for BL development(10, 11), including mutations in *TP53*, *ID3, TCF3,* and *CCND3*(12–14), activated phosphatidylinositol 3-kinase (PI3K) signaling pathway(15) and inactivation of tumor suppressors like *ARF1*(16) and *DDX3X*(17). However, the precise combinations of altered activated and repressed transcriptional pathways in BL have not been fully explored. Differential mRNA expression and complementary miRNA profiling studies can be rich in biological information, since variations in expression of large numbers of protein-coding genes can be captured by miRNAs that regulate them. miRNAs are single-stranded, non-coding ribonucleic acid molecules encoded in the genome with an average length of 19 to 22 nucleotides. They modulate mRNA expression post-transcriptionally through base-pairing to complementary sequences at the 3’-UTR of numerous target mRNAs, leading to mRNA translational repression or degradation(18). The impact of mRNA-miRNA interactions propagates along large gene expression networks by targeting of numerous specific transcripts and regulatory elements such as transcription factors. Emerging evidence suggests that abnormal modulation of mRNA transcription via miRNAs might be a significant factor in BL pathogenesis(19).

The aim of this pilot study was to analyze mRNA and miRNA expression profiles in pediatric BuAL and BuLY. An evaluation of aberrant mRNA and miRNA expression changes in BuAL and BuLY compared to its normal counterpart could provide an insight into mechanisms involved in Burkitt lymphomagenesis and identify key underlying miRNAs and miRNA-mRNA interactions. A further aim of the study was to analyze transcriptomic differences and miRNA-mRNA networks that distinguish BuAL from BuLY subtypes.

## Methods

### Patient samples

This study was approved by the Ethics Committees of the University of Rostock (Reference number A 2023-0054) and the Hospital Sant Joan de Déu, Barcelona (Reference number HSJD C.I. PIC-36-23). Specimen collection was conducted in accordance with the ethics guidelines for the use of human material and in accordance with the Declaration of Helsinki. Informed consent was obtained from all parents or legal guardians.

Total RNA from 13 fresh tumor tissue samples of pediatric BuAL and BuLY and a control group composed by a pool of four normal control lymphoid tissue (NCLT) mixes were included in the study.

### Clinical characteristics overview

BL cases included in this pilot study (BuAL n=7, BuLY n=6) were diagnosed between 2009-2021 at the Hospital Sant Joan de Déu of Barcelona, Spain, all fulfilling the diagnostic criteria for BuAL/BuLY according to the International Consensus Classification of Mature Lymphoid Neoplasms(3) and the 5^th^ edition of the WHO-Classification on mature lymphoid neoplasms(5). BuAL was defined as more than 25% blasts in BM by cytomorphology and was confirmed by flow cytometry (range 32-100%, median 38%). Cases of precursor-B-cell acute lymphoblastic leukemia with *IG::MYC*(*20*), which is considered a different entity, were excluded from this study. Hematoxylin and eosin-stained slides of the selected paraffin blocks were examined by experienced hematopathologists to ensure sample sufficiency and quality. All biopsy samples were Epstein-Barr virus-encoded small RNA (EBER) negative. *MYC* rearrangement was positive by fluorescent *in situ* hybridization in all cases. The clinical characteristics are summarized in supplementary file 1. The median age at diagnosis was 11 years (range 2-18 years), 77% of patients were male and 23% were female. All cases were *de novo* and HIV negative. Cases were homogeneously treated with the Inter-B-Ritux-2010 Protocol(7) starting in 2010, one case before 2010 received treatment with the LNH-B-04 protocol. After flow cytometric analysis showing no evidence of malignant cell infiltration, exceeding material from normal tonsils (n=2) and a normal lymph node (n=1) from pediatric patients (median age 9 years) was used to create NCLT mixes for comparison with BL.

### 2. Total RNA isolation

A total of 17 samples were included, 13 fresh tumor tissue samples of BuAL/BuLY cases and four NCLT mixes. Fresh BuAL bone marrow samples underwent treatment with a Ficoll-Hypaque (Sigma, St Louis, MO, USA) centrifugation gradient prior to RNA extraction. Total RNA from fresh BuLY homogenates, BuAL and normal lymphoid tissue was extracted using TRIzol using TriPure Isolation Reagent (Roche Diagnostics, Indianapolis, IN, USA) and Direct-zol RNA MiniPrep kit (Zymo Research, California, USA) and stored at –80°C. In addition, two Burkitt lymphoma cell lines, Raji and DG-75, were sequenced as external reference samples to contextualize the global expression profiles in the principal component analysis. The cell lines were excluded from differential expression analysis because they were not considered biologically comparable replicates of the primary samples.

For mRNA/miRNA extraction from cell lines Raji and DG-75 the miRNeasy Micro Kit (Qiagen, Hilden, Germany), was used as per manufactureŕs instructions. Qubit (Thermo Fisher Scientific, Waltham, USA) was used for RNA quantification and Agilent Bioanalyzer 2100 (Agilent Technologies, CA, USA) was employed to evaluate the RNA integrity. Mean RNA integrity (RIN) value of samples was 7.5 (range 4.9-9.2) (supplementary file 2, supplementary methods).

### 3. Transcriptome sequencing

The mRNA-sequencing service was provided by Novogene® (Beijing, China). In brief, mRNA purification was performed using poly-T-oligo-attached magnetic beads. Reverse-transcription to cDNA was performed, which was then processed for sequencing library preparation. After library quality control, Illumina sequencing was performed generating fastq files. Data quality control was carried out prior to bioinformatic alignment and analysis (supplementary file 2, supplementary methods and supplementary figures). Data can be accessed at the GSE337108 accession number.

### 4. miRNA profiling

miRNA expression analysis was evaluated using the NanoString nCounter System Human v3 miRNA assay (Nanostring, Seattle, WA, USA) (supplementary methods). miRNA profiling of eleven cases of this series was part of a previous publication(21). The datasets generated and analyzed are available in the NCBI’s Gene Expression Omnibus, accession number GSE 337806.

### 5. Bioinformatic analysis

#### Differential gene expression analysis

DE was performed with DESeq2 pipeline using standard pairwise contrasts, and protein-coding genes meeting both false discovery rate (FDR)<0.05 and |log₂ fold change(FC)|>0.5 criteria, were considered as differentially expressed. Direction-specific DEG lists (up/down per contrast) derived from this pipeline were visualized for overlap using the ggvenn R package (0.1.19), and customized volcano plots were generated with the EnhancedVolcano R package (1.24.0) to highlight representative hits. For visual clarity, volcano plots applied internal cutoffs of raw p< 0.01 and |log₂FC|> 1 for labeling of genes with the strongest effect size.

#### Gene Ontology (GO) and Kyoto Encyclopedia of Genes and Genomes (KEGG) enrichment analysis of DEG

GO enrichment analysis of DEG was implemented by the clusterProfiler R package, in which gene length bias was corrected. GO terms with corrected p-value less than 0.05 were considered significantly enriched by DEG. KEGG is a database resource for understanding high-level functions and utilities of the biological system, such as the cell, the organism and the ecosystem, from molecular-level information, especially large-scale molecular datasets generated by genome sequencing and other high-through put experimental technologies (http://www.genome.jp/kegg/)(22–24). We used clusterProfiler R package to test the statistical enrichment of differential expression genes in KEGG pathways. Gene Set Enrichment Analysis (GSEA) is a computational approach to determine if a predefined gene set can show a significant consistent difference between two biological states. The genes were ranked according to the degree of DE in the two samples, and then the predefined gene sets were tested to see if they were enriched at the top or bottom of the list. GSEA can include subtle expression changes. We used the local version of the GSEA analysis tool http://www.broadinstitute.org/gsea/index.jsp. GO, KEGG data set were used for GSEA independently.

#### Functional Enrichment Analysis

To interpret the biological significance of the differential gene expression profiles, we performed over-representation analyses (ORA) on the sets of up– and down-regulated genes obtained from each main contrast (BL vs NCLT and BuAL vs BuLY). ORA was carried out using the clusterProfiler (4.14.0) R package(25), implementing the compareCluster function with the enrichPathway routine from the ReactomePA (1.50.0) package to map genes onto curated REACTOME pathways. This approach, applied independently to the four direction-specific DEG lists, enables a more resolved biological interpretation by distinguishing pathways associated with transcriptional activation from those linked to repression. Significance thresholds were set at adjusted *p*-value<0.05 (Benjamini–Hochberg (BH) correction). In parallel, to complement the Reactome results and capture broader biological programs, Gene Ontology Biological Process (GO-BP) enrichment was also performed using clusterProfiler R package (4.14.0) followed by semantic similarity clustering with the simplifyEnrichment (2.0.0) package. This framework groups redundant GO terms into coherent functional clusters based on pairwise similarity metrics, generating a consolidated “meta-enrichment” map that highlights recurrent biological themes across contrasts while minimizing interpretational redundancy(26).

#### Gene set enrichment analysis (GSEA)

Finally, to further assess coordinated, rank-based transcriptional programs beyond discrete DEG thresholds, GSEA was conducted using the MSigDB Hallmark collection(27), retrieved programmatically via the msigdbr R package (25.1.1). Hallmark gene sets were filtered to include 15– 500 members and only sets with ≥5 overlapping genes with the input ranking were evaluated. Enrichment scoring was carried out within the decoupleR framework (2.12.0)(28), using a signed gene-level statistic defined as sign (log₂FC) × (−log₁₀ FDR) derived per contrast. To improve robustness, multivariate linear modeling was applied to account for shared gene membership across sets. Results were summarized as normalized enrichment scores (NES) with BH-adjusted p-values.

#### Pathway & Transcriptional Factor Activity Analysis

To estimate the direction and relative activity of cancer-relevant signaling pathways, we applied the PROGENy resource method (Pathway RespOnsive GENes)(29) in combination with the decoupleR R package (2.12.0) statistical framework(28), implementing the multivariate linear model algorithm. PROGENy infers pathway activity from gene expression data using experimentally derived perturbation signatures that link transcriptomic responses to upstream pathway perturbations. In this analysis, normalized pathway activity scores were computed for 14 canonical signaling pathways using the top 500 most responsive genes per pathway and a signed statistic defined as sign(log₂FC) × (−log₁₀ FDR) derived from each contrast. The resulting NES reflect the relative activation or repression of each pathway between conditions, allowing direct comparison of pathway-level dynamics across the two biological contrasts.

Next, transcription factor (TF) regulatory activities were inferred using DoRothEA regulons from the dorothea R package (1.18.0), a curated resource of signed TF–target interactions stratified by confidence levels. For this study, only high-confidence regulons (A–C) were retained(30). TF activity estimation was performed within the decoupleR framework (2.12.0) using the univariate linear model, which provides stable activity estimates in the presence of overlapping regulons. As input, we supplied the same signed statistics used for the PROGENy pathway analysis (sign(log₂FC) × (−log₁₀ FDR)) derived per contrast, and TFs were evaluated only when their corresponding regulons met a minimum overlap of ≥20 target genes in the expression set. Multiple-testing correction across TFs was applied using the BH procedure.

#### miRNA profiling

Raw data from the nCounter Digital Analyzer were imported to the Rosalind® software (https://rosalind.bio/) for analysis. Standard settings for background subtraction and normalization were applied (supplementary methods). Group-wise comparisons were performed with the t-test method, delivering FC and p-values, with p-value adjustment through the BH method of estimating FDR.

#### miRNA-mRNA integrative ranking analysis

miRNA–mRNA integrative analysis was conducted by combining DE, inverse expression correlation, and experimentally validated miRNA–target evidence. Differentially expressed miRNAs were defined from the provided miRNA DE tables using FDR<0.05 and |FC|>1.5, while protein-coding DE mRNAs were defined from DESeq2 pairwise contrasts using FDR<0.05 and |log₂FC|> 0.5 (as stated above). For each DE contrast independently (BL vs NCLT and BuAL vs BuLY), miRNA normalized expression and mRNA variance stabilizing transformation expression (blind=TRUE) were aligned by sample, and Spearman rank correlations were computed across the subset of samples belonging to the corresponding contrast group, using the Hmisc R package (5.2-4). Multiple testing across all tested miRNA–mRNA pairs within each contrast was controlled using the BH procedure, and only significant inverse associations were retained (ρ≤−0.5 and BH-adjusted correlation p-value<0.05). To prioritize biologically supported interactions, inverse-significant correlation pairs were intersected with experimentally validated miRNA–target relationships retrieved via the multiMiR R package (1.28.0; database version 2.4.0, integrating “validated” evidence from miRecords, miRTarBase, and TarBase). Multi-miRNA probe identifiers containing “+” were split into individual miRNA components for database querying and then collapsed back to the original probe label while retaining component-level evidence. For each retained validated miRNA–mRNA interaction (“edge”), evidence support was summarized (number of supporting databases and unique PubMed identifiers), and interactions were annotated with miRNA and mRNA DE statistics (FC/log₂FC and FDR) to flag expected regulatory directionality (opposite sign between miRNA and target mRNA effects). Final results were generated per contrast as an edge-level table of validated, inverse-significant miRNA–mRNA interactions and a miRNA-level summary table ranked by the number of validated DE targets observed among inverse-significant correlations, with correlation and evidence statistics retained as supporting metadata. Gene enrichment and functional annotation analysis of the up– and downregulated target transcripts was performed with the DAVID online tool(31). GO terms and annotation categories were selected with the threshold of significance being defined as *p*<0.01 and FDR<0.05.

## Results

### Differential mRNA expression in Burkitt leukemia and lymphoma

Aiming to disentangle the transcriptional programs that distinguish BuAL from BuLY and from normal lymphoid tissue, we prioritized two main biological comparisons: (a) BL (BuAL and BuLY combined) versus (vs) NCLT and (b) BuAL vs BuLY, comparing the two Burkitt subtypes directly. After de-duplication to one symbol per gene and restriction to protein-coding transcripts, DEG were defined using a significance threshold of an adjusted *p*-value (FDR< 0.05) and |log₂FC|> 0.5.

In the BL vs NCLT contrast, a total of 1965 DEGs were identified, including 1214 up-regulated and 751 down-regulated genes, indicating broad transcriptional remodeling between tumor and normal tissue (Figure 1a). Strongly up-regulated genes in the BL vs NCLT contrast included *DEFA1B, GABRA3, APOC3, GRM6, and HOXB1*, several of which have been implicated in innate immune responses(32), regulation of transcription(33) and self-renewal of hematopoietic progenitor cells(34). Conversely, down-regulated genes such as *CCL17, SIM1, CRH, and CSF2* are involved in cytokine (35) and immune signaling(36), pointing to an attenuation of inflammatory and chemotactic signaling in BL samples (Figure 1a). In contrast, the BuAL versus BuLY comparison revealed 4519 DEGs, with a pronounced predominance of down-regulated genes in BuAL, 4143 relative to 376 up-regulated DEGs, suggesting extensive transcriptional repression in BuAL compared with BuLY (Figure 1b). In the BuAL versus BuLY comparison, the most prominent up-regulated genes were *DEFA1, DEFA1B, DEFA3, DEFA4, and BEX1*, implicated in innate immune response pathways (37), proliferation and invasiveness of malignant cells in the leukemic context(38). In contrast, *POTEM, MYOM3, TNFRSF11B*, *PRSS38*, and *CALB2* were among the most strongly down-regulated genes, several of which are linked to adhesion, bone-remodeling(39) and neuronal-like signaling processes in cancer(40)(Figure 1b).

**Figure 1.**
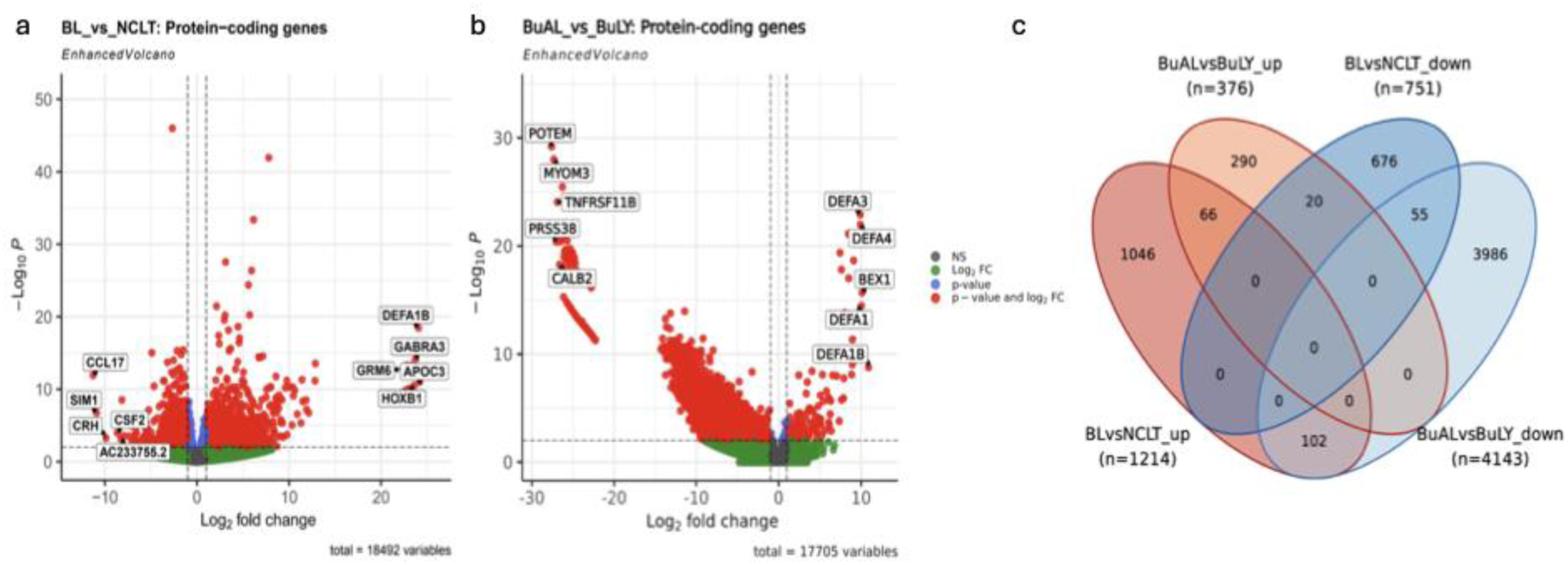
a_DEG in BL vs NCLT and b_DEG in BuAL vs BuLY. Volcano plots generated with the EnhancedVolcano R package (v1.24.0), displaying differential expression of protein-coding genes (n = 18492) between BL samples and NCLT. DEG were defined using the DESeq2 pipeline applying statistical thresholds of FDR<0.05 and |log₂FC|>0.5. From this set of significant DEGs, the top five most upregulated and top five most downregulated transcripts were selected based on absolute log₂FC and annotated on the plot. For visualization purposes, the volcano plot displays log₂FC on the x-axis and – log₁₀(raw p-value) on the y-axis. Genes meeting the following cutoffs (raw p-value< 0.01 and |log₂FC|> 1) are colored in red. Dashed vertical and horizontal reference lines indicate log₂FC = ±1 and raw p-value = 0.01, respectively; these serve only as graphical guides to highlight high-magnitude changes and do **not** reflect the statistical cutoffs used to define DEGs. **c_Overlap of direction-specific DEG sets across contrasts.** Four protein-coding DEG lists (DESeq2 DE pipeline; FDR<0.05 and |log₂FC|> 0.5) from the main 2 comparisons (BL vs NCLT & BuAL vs BuLY) were compared with the ggvenn R package (0.1.19): BLvsNCLT_up, BLvsNCLT_down, BuALvsBuLY_up, and BuALvsBuLY_down. Each ellipse corresponds to one list; labels on the plot give the size of each set and of every intersection. The diagram shows that most DEGs are list-specific (e.g., large unique sectors for BLvsNCLT_up and BuALvsBuLY_down), with limited shared signal: a modest up-up overlap (66 genes) and down-down overlap (55 genes), and small cross-direction overlaps (BLvsNCLT_up with BuALvsBuLY_down = 102 common genes; BuALvsBuLY_up with BLvsNCLT_down = 20 genes). Overall, the pattern indicates largely context-specific transcriptional programs with only focal commonalities between contrasts.

A four-way Venn analysis (Figure 1c) further clarified the relationships between the two contrasts. While the majority of genes were uniquely regulated within each comparison, several partial overlaps emerged: 66 genes were jointly up-regulated in both contrasts, 55 genes were consistently down-regulated, 102 genes were up in BL vs NCLT but down in BuAL vs BuLY, and a smaller subset of approximately 20 genes showed the inverse pattern (down in BL vs NCLT, up in BuAL vs BuLY) (supplementary file 3). These limited yet biologically coherent intersections suggest that a minor fraction of pathways may contribute to both malignant transformation and phenotypic divergence between leukemia and lymphoma. The overall scarcity of shared DEGs, however, reinforces the notion that the tumor versus normal and subtype axes capture largely independent layers of transcriptional regulation. Overall, these results indicate that while the BL vs NCLT comparison captures canonical tumor–normal transcriptional remodeling, the BuAL vs BuLY axis reveals a distinct, predominantly repressive landscape specific to the leukemic phenotype. Together, these findings delineate complementary transcriptomic dimensions of BL, setting the stage for the functional and pathway-level analyses described below.

### Mapping gene expression programs to functional networks

#### Functional enrichment of direction-specific gene sets reveals distinct proliferative, immune, and structural modules

To delineate the molecular programs driving the observed transcriptional differences, we performed functional enrichment analyses by mapping the up– and down-regulated DEGs from each main contrast (BL vs NCLT and BuAL vs BuLY) to curated REACTOME pathways. This approach, using four direction-specific gene lists in total, enables finer biological resolution by separating processes linked to transcriptional activation from those linked to repression, while allowing direct cross-comparison across the two biological contrasts (see materials and methods).

In the BL vs NCLT comparison, the upregulated subset (n = 1214) showed strong enrichment for cell-cycle progression, mitotic checkpoint regulation, and mitochondrial translation pathways, reflecting the enhanced proliferative and biosynthetic activity typical of tumor tissue relative to normal controls. Conversely, the downregulated genes (n = 751) were predominantly involved in immune and cytokine-related signaling, including Interleukin-2/6/20 family signaling and T-cell costimulation by CD28, consistent with the suppression of cytokine-mediated immune communication within the tumor microenvironment (Figure 2a). In the BuAL vs BuLY comparison, the upregulated subset (n = 376) was characterized by a distinct enrichment in innate and antimicrobial defense pathways, notably neutrophil degranulation, antimicrobial peptides, and α-defensins. In contrast, the downregulated genes (n = 4143) were markedly enriched for extracellular matrix (ECM) organization, ECM proteoglycans, cell-adhesion, and neuronal system modules, indicating a loss of adhesion and microenvironmental structure in leukemia relative to lymphoma (Figure 2a). Taken together, the Reactome results highlight sharply divergent biological signatures between the two disease contexts: BL vs NCLT dominated by proliferative and metabolic activation accompanied by immune suppression, and BuAL vs BuLY defined by innate immune stimulation alongside the erosion of adhesion and ECM integrity. To complement the Reactome analysis, we next performed GO-Biological Process (GO-BP) enrichment using a robust clustering framework that consolidates semantically related terms into higher-order biological modules (see materials and methods). The GO-BP enrichment map revealed coherent, biologically interpretable clusters, whose distribution across the four DEG sets paralleled the Reactome enrichment findings. Specifically, cell-cycle and metabolic clusters were driven almost exclusively by BL vs NCLT upregulated genes, whereas interleukin and immune-response modules were associated with BL vs NCLT downregulated genes (Figure 2b). On the contrary, in the leukemia-versus-lymphoma contrast, ECM and adhesion clusters, together with developmental and neuronal-related processes, were mainly contributed by BuAL vs BuLY downregulated genes, while defense and inflammatory response modules aligned with BuAL vs BuLY upregulated genes. Overall, the GO-BP functional analysis confirmed that the two major biological transitions involve fundamentally distinct functional rewiring: a proliferative and biosynthetic shift in tumor versus normal tissue, and an immune/structural reorganization distinguishing the leukemic from lymphomatous BL subtypes (Figure 2b).

**Figure 2.**
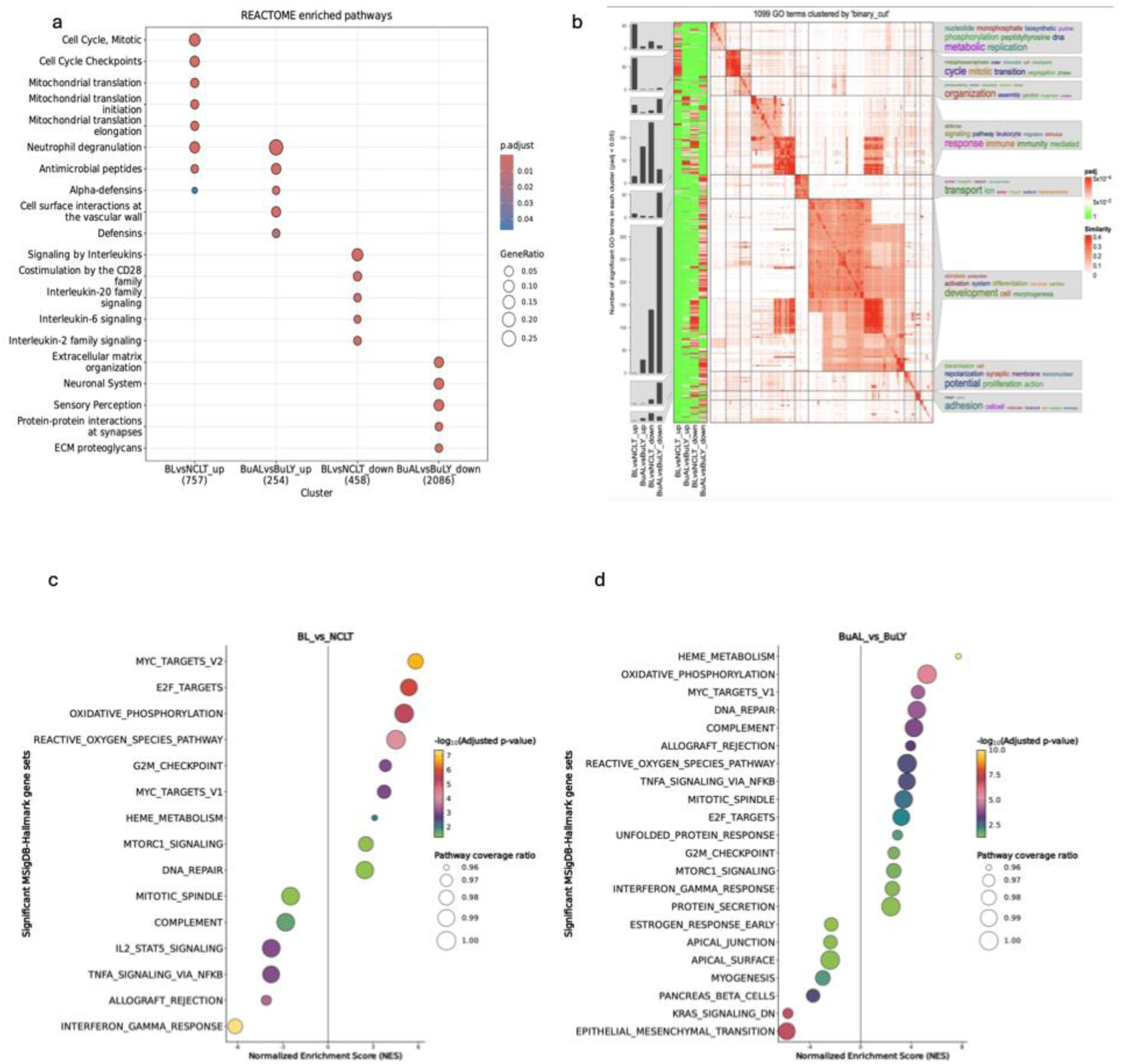
a_REACTOME pathway enrichment across direction-specific gene sets in Burkitt lymphoma and leukemia. Dot plot summarizing over-representation analysis (ORA) results obtained with clusterProfiler (4.14.0) and ReactomePA (1.50.0) R packages using four direction-specific DEG lists: BLvsNCLT_up, BLvsNCLT_down, BuALvsBuLY_up, and BuALvsBuLY_down. Each dot represents a significantly enriched Reactome pathway; its color encodes adjusted p-value, and size denotes the proportion of genes within that pathway relative to the input list (GeneRatio). Pathways are ordered by significance, and gene counts for each input set are shown in parentheses on the x-axis. **b_Functional clustering of GO-BP terms across Burkitt transcriptomic contrasts.** Heatmap-based functional map generated using the simplifyEnrichment R package (2.0.0) framework, applied to GO-BP over-representation results from the four direction-specific DEG lists (BLvsNCLT_up, BLvsNCLT_down, BuALvsBuLY_up, BuALvsBuLY_down). GO terms were clustered according to semantic similarity, producing meta-clusters that group biologically related processes. Each cell in the similarity matrix reflects pairwise overlap between GO terms (red = higher similarity). The side barplot indicates the number of significant GO terms per cluster, while the vertical annotation panel shows their distribution across the four DEG sets (green = significant enrichment). Representative word clouds summarize the most frequent biological themes within each functional cluster. Adjusted p-values are color-scaled also from green to red to denote enrichment strength. **c, d_Hallmark gene set enrichment across direction-specific gene sets in BL and NCLT and BuAL vs BuLY.** Dot plot summarizing normalized enrichment-scores obtained with msigdbr R package (25.1.1), using two DEG lists: BLvsNCLT and BuALvsBuLY. Each dot represents a significantly enriched Reactome pathway; its color encodes adjusted p-value, and size denotes the proportion of genes within that pathway relative to the input list (GeneRatio). Pathways are ordered by significance, and gene counts for each input set are shown in parentheses on the x-axis.

Finally, to assess coordinated, rank-based transcriptional programs beyond discrete DEG thresholds regarding enrichment in hallmark gene sets, GSEA was conducted using the MSigDB Hallmark collection (see materials and methods). Regarding the BL vs NCLT contrast, MYC, E2F, oxidative phosphorylation and reactive O2 species signaling emerged as the most strongly activated pathways, concordant with a drive to progress from G1 to S phase, as well as a metabolic dependency on oxidative phosphorylation (Figure 2c). Conversely, repressed IFN-gamma response, TNF-alfa signaling via NFkB and IL2 mediated STAT5 signaling were observed, consistent with attenuated Th1-immune response, decreased antiapoptotic effect and immune evasion relative to NCLT. In the BuAL vs BuLY comparison, several cascades displayed concordant behavior across both contrasts with reinforcement in the leukemic state: MYC, E2F, oxidative phosphorylation and reactive O2 species signaling are consistently activated, underscoring their central role in Burkitt leukemia and lymphoma (Figure 2d). This was accompanied by repression of the epithelial-mesenchymal transition in Burkitt leukemia compared to Burkitt lymphoma, consistent with reduced homing in the bone marrow of the leukemic form. The IFN-gamma signaling pathway and TNF-alpha signaling via NFkB exhibited opposite directionality, being repressed in BL vs NCLT but activated in BuAL vs BuLY, highlighting a switch from reduced apoptotic effect and reduced immunosurveillance through the IFN-gamma pathway in BL vs NCLT and its reinforcement, linked to increased JAK-STAT signaling observed in the leukemic state.

#### Transcription Factor Activity Pathway and transcriptional factor activity modeling reveals shared and divergent signaling dynamics across Burkitt leukemia and lymphoma subtypes

To further evaluate the transcriptional differences at the TF signaling level, TF activity estimation was performed within the decoupleR framework (see materials and methods). Regarding the BL vs NCLT contrast, a MYC-driven proliferative backbone emerged as the most strongly expressed TF signaling pathway, coupled with E2F4 and MYB pathway activity. Conversely, the TF signaling pathways of SPIB, RBPJ and TBX21 are repressed, consistent with impaired differentiation, repressed Notch-signaling and decreased Th1-immune response relative to normal tissue. For the Burkitt leukemia versus lymphoma contrast, a common MYC-driven proliferative backbone, coupled with E2F4 and MYB pathway activity was observed, as well as Burkitt leukemia specific enrichment in SPI1, LYL1, NRF1 and ZFX transcription factor signaling activity, TF that are coupled with hematopoietic self-renewal(41) and differentiation(42) (Figures 3a and 3b).

**Figure 3.**
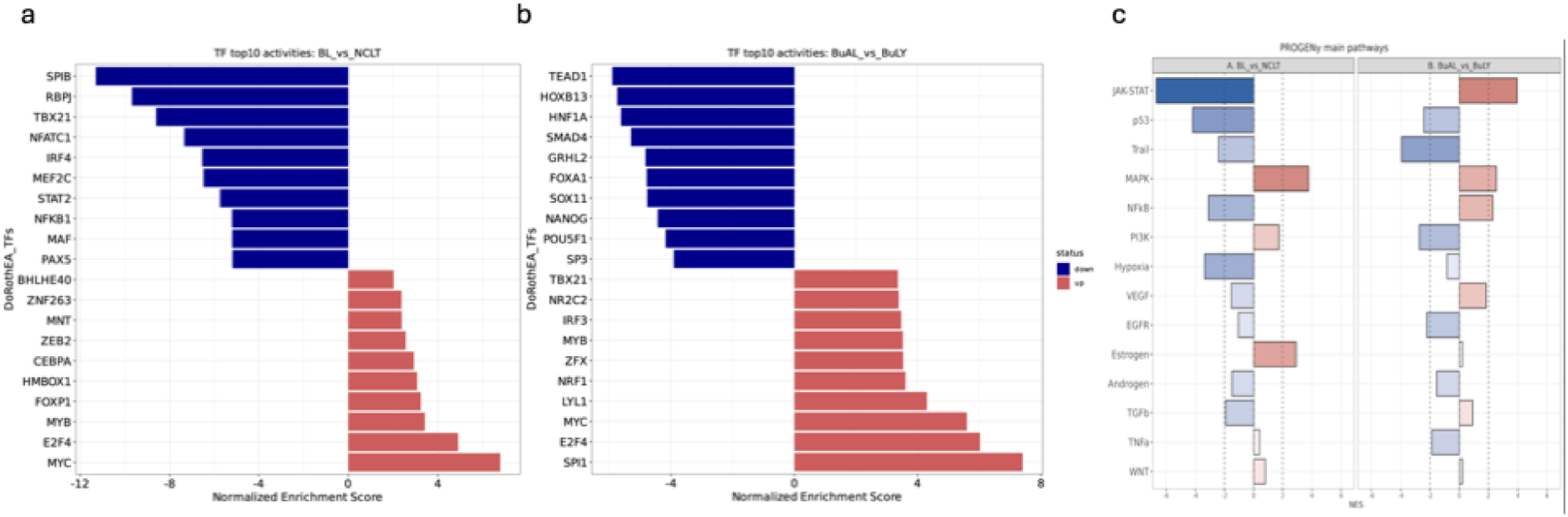
a and b_ Differential TF activities in BL vs NCLT and in BuAL vs BuLY inferred by DoRothEA. Barplots display NES of 20 TF signaling pathways estimated from transcriptomic profiles using the dorothea R package (1.18.0). Panels represent the two principal biological contrasts: **a_ BL vs NCLT** and **b_ BuAL vs BuLY**. Positive and negative NES values indicate predicted activation or repression of the corresponding pathways, respectively. Blue and red color gradients denote the direction and magnitude of pathway activity changes. **c_Differential signaling pathway activities in BL inferred by PROGENy**. Barplots display NES of 14 canonical signaling pathways estimated from transcriptomic profiles using the PROGENy-decoupleR computational model (https://saezlab.github.io/decoupleR/articles/pw_bk.html). Panels represent the two principal biological contrasts: a_ BL_vs_NCLT and b_ BuAL_vs_BuLY. Positive and negative NES values indicate predicted activation or repression of the corresponding pathways, respectively. Blue and red color gradients denote the direction and magnitude of pathway activity changes. Vertical dotted lines (at +2 and −2) mark the empirical threshold for strongly deregulated pathways.

To further characterize the transcriptional differences at the signaling level, we estimated the activities of 14 critical signaling pathways using the PROGENy framework, which infers upstream pathway perturbations from downstream transcriptional footprints (see materials and methods). Regarding the BL vs NCLT contrast, MAPK, Estrogen-receptor and PI3K signaling emerged as the most strongly activated pathways, consistent with the proliferative and biosynthetic drive typical of malignant transformation. Conversely, JAK–STAT, p53, NFkB, hypoxia and TRAIL pathways were repressed, suggesting attenuated cytokine, apoptotic, and stress-response signaling relative to normal lymphoid tissue (Figure 3c). In the BuAL vs BuLY comparison, pathway activation shifted toward JAK– STAT, NF-κB, MAPK, TGF-β, and VEGF signaling, reflecting heightened inflammatory, angiogenic, and cytokine-related activity in the leukemic phenotype. This was accompanied by repression of TP53, TRAIL, EGFR, and PI3K pathways, consistent with impaired apoptotic control and altered growth-factor signaling in Burkitt leukemia relative to Burkitt lymphoma (Figure 3c). Notably, several cascades displayed concordant behavior across both contrasts: MAPK signaling was consistently activated, underscoring its central role in sustaining proliferative signaling, whereas TP53 and TRAIL pathways were downregulated in both, indicating shared attenuation of apoptosis-related responses across BuAL and BuLY. In contrast, JAK–STAT signaling exhibited opposite directionality, being repressed in BL vs NCLT but activated in BuAL vs BuLY, highlighting a switch from reduced cytokine signaling through the JAK-STAT pathway in BL versus normal and its reinforcement in the leukemic state.

#### miRNA-mRNA integrative analysis reveals distinct proliferative, immune, and structural modules

The group-wise comparison of the miRNA expression profiles obtained by the NanoString nCounter assays revealed the following results:

#### Differentially expressed miRNAs

Expression profiling analysis of BL compared to NCLT showed eleven miRNAs which were differentially expressed (FC>1.5, *p*-adj<0.05). Three miRNAs were differentially upregulated in BL compared to NCLT and eight miRNAs were downregulated (supplementary table 1). The higher expressed miRNAs in BL compared to NCLT were hsa-miR-106a-5p, hsa-miR-17-5p and hsa-miR-7-5p. The lower expressed miRNAs in BL compared to NCLT included hsa-miR-150-5p, hsa-miR-450b-5p and hsa-miR-342-3p, among others (Figure 4a). Expression profiling analysis comparing BuAL (n=7) with BuLY (n=6) showed 661 miRNAs DE in BuAL compared with BuLY with FC>1.5, *p-*adj<0.05, 30 upregulated and 631 downregulated (supplementary table 2). The higher expressed miRNAs in BuAL compared to BuLY were, among others, hsa-miR-223-3p, hsa-miR-144-3p, hsa-miR-142-3p, hsa-miR-150-5p and hsa-miR-15a-5p, whereas the lower expressed miRNAs in BuAL compared to BuLY included, among others, hsa-miR-1285-5p, hsa-miR-1972, and hsa-miR-4787-3p among others (Figure 4b).

**Figure 4.**
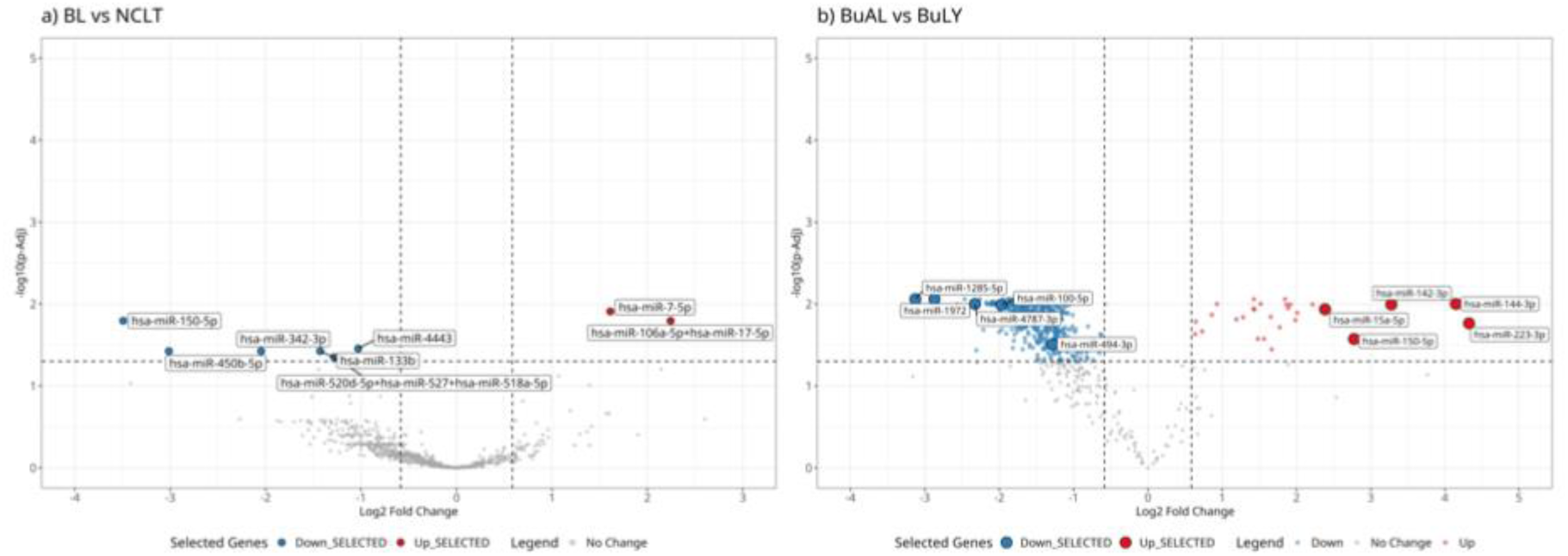
a_Differential miRNA expression in BL versus NCLT (FC>1.5, p-adj<0.05). b_ Differential miRNA expression in BuAL versus BuLY (FC>1.5, p-adj<0.05). Volcano plots: X-axis log_2_FC, Y-axis log_10_ p-adj. Volcano plot: Differential miRNA expression intensity: red color tones upregulation, blue color tones downregulation.

### miRNA and mRNA network analysis

#### 1. BL versus NCLT

Inverse-significant miRNA–mRNA interactions ranked by the number of validated DE targets observed among inverse-significant correlations, with correlation and evidence statistics retained as supporting metadata in BL vs NCLT are shown in table 1. A total of 244 edge-level validated, inverse-significant miRNA–mRNA interactions were identified, with 46 upregulated and 198 downregulated validated targets (supplementary file 4). Visualized data in the form of a miRNA-target network (STRING version 12.0) are shown in Figure 5a. Functional enrichment analysis of the inversely-expressed edge-level validated target genes of the DE miRNAs provided us with an overall clue of their functional roles in BL, showing that whereas the 46 upregulated validated targets of DE downregulated miRNAs are significantly associated with genes enriched in activation of transcription, nucleic acid binding and nucleus and nucleoplasm organization pathways (Figure 5b), the 198 downregulated validated targets are significantly associated with pathways of immune system processes and T-lymphocyte activation (FDR<0.05) (Figure 5c) (supplementary files 5 and 6). Transcription factor pathways including MYC-MAX, NMYC, E2F, FOXO and YY1 among others, showed enrichment by both up-and downregulated mRNA targets of DE miRNAs, highlighting a fine-tuning of TF pathways in BL by DE miRNAs (supplementary files 5 and 6).

**Figure 5.**
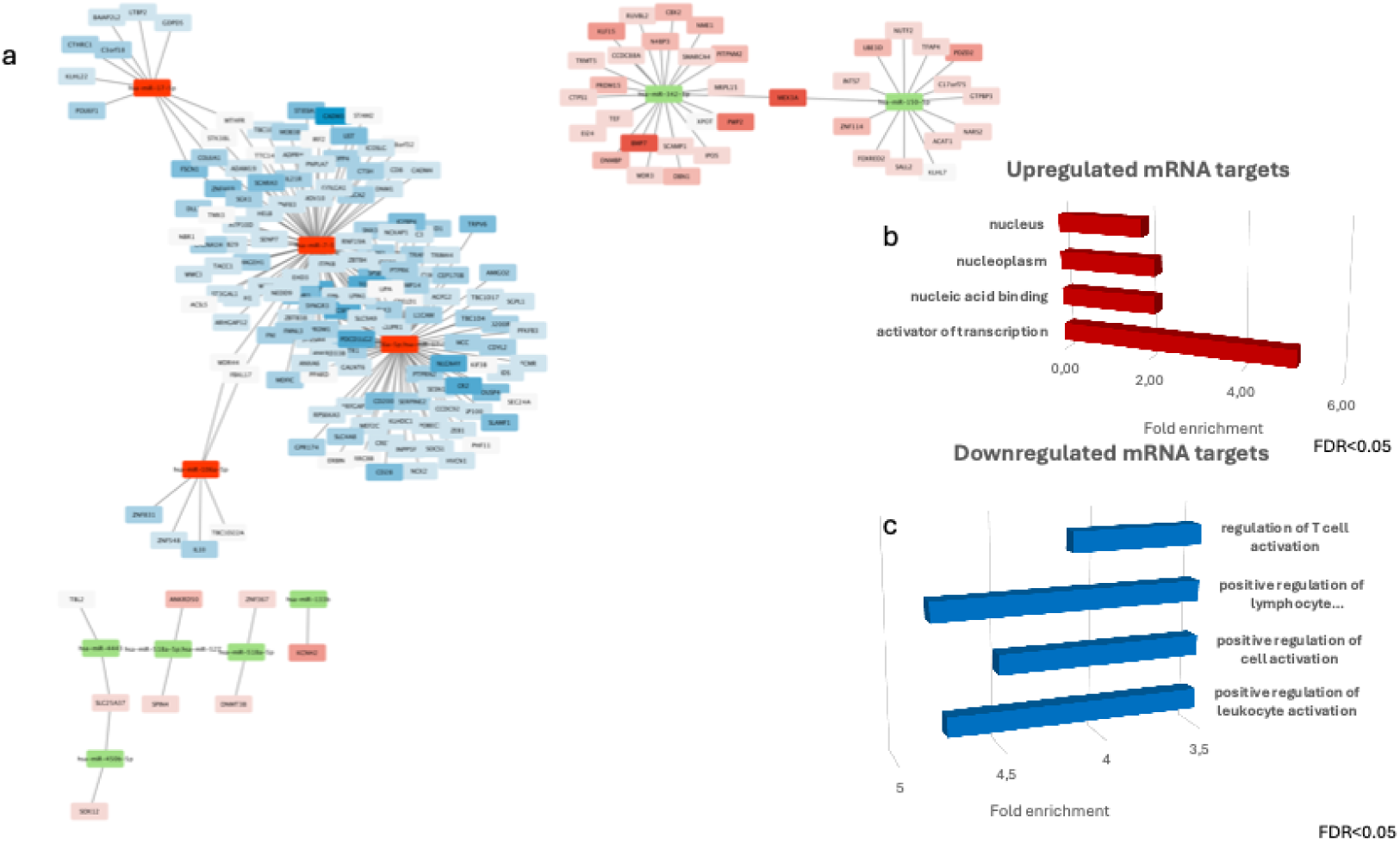
a_miRNA-mRNA target network analysis in BL versus NCLT performed with STRING (version 12.0) including DEmiRNAs (FC>1.5; p-adj<0.05) and their validated targets according to the multiMiR R package; log_2_FC>|0.5|; p-adj<0.05). Only significant inverse associations were retained (ρ ≤ −0.5 and BH-adjusted correlation p-value<0.05). Upregulated miRNAs in red color, downregulated miRNAs in green color. Upregulated validated mRNA targets with significant edges in red color tones, downregulated validated mRNA targets with significant edges in blue color tones. **b**_ **DAVID Functional enrichment analysis** of **upregulated targets** (red color tones) fulfilling edge-level validated, inverse-significant miRNA–mRNA interactions (FDR<0.05). **c**_ **DAVID Functional enrichment analysis of downregulated targets** (blue color tones) fulfilling edge-level validated, inverse-significant miRNA– mRNA interactions (FDR<0.05).

**Table 1.**
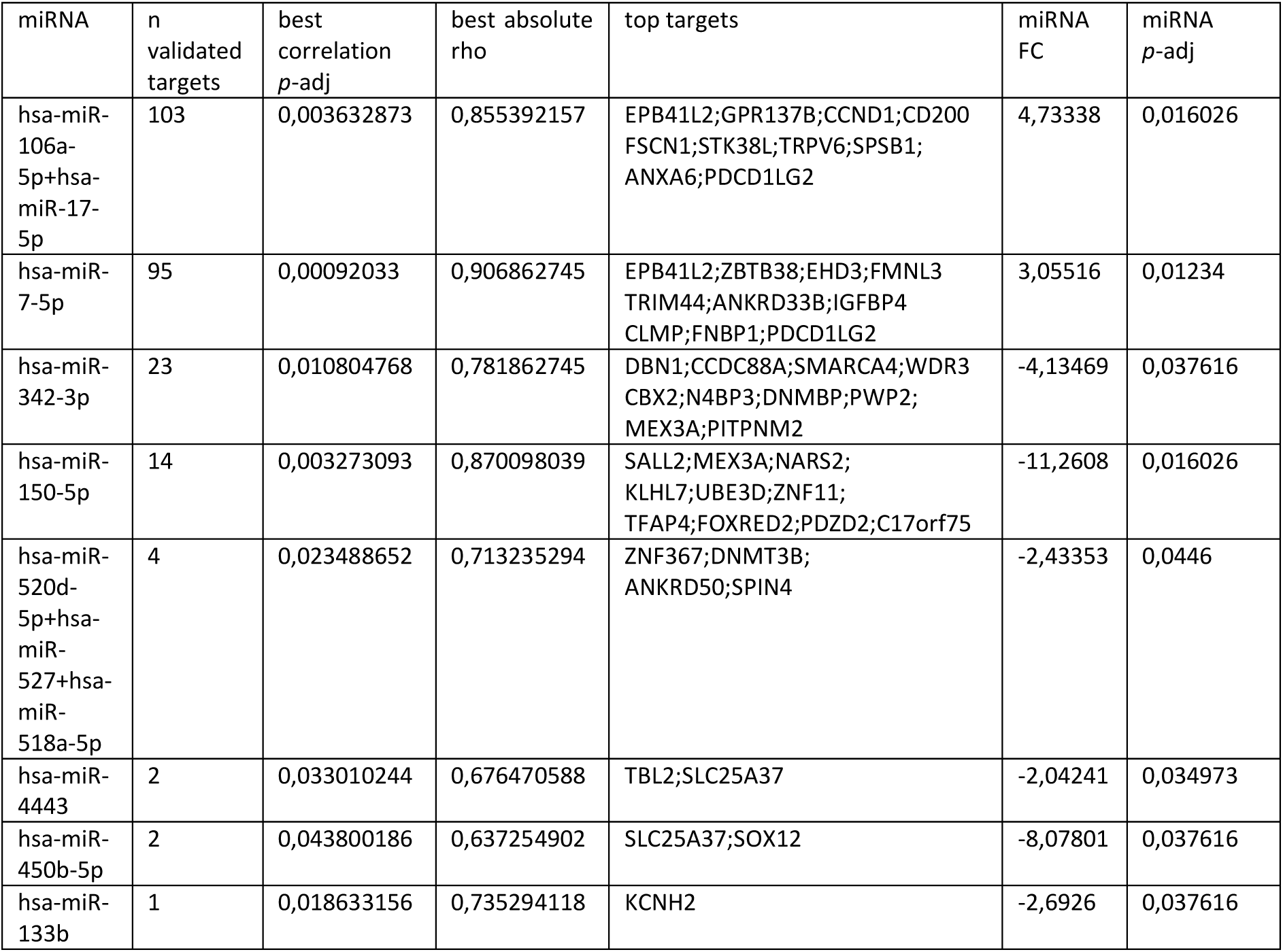
Inverse-significant miRNA-mRNA interactions ranked by number of validated DE targets observed in BL versus NCLT.

| miRNA | n validated targets | best correlation *p*-adj | best absolute rho | top targets | miRNA FC | miRNA *p*-adj |
| --- | --- | --- | --- | --- | --- | --- |
| hsa-miR-106a-5p+hsa-miR-17-5p | 103 | 0,003632873 | 0,855392157 | EPB41L2;GPR137B;CCND1;CD200 FSCN1;STK38L;TRPV6;SPSB1; ANXA6;PDCD1LG2 | 4,73338 | 0,016026 |
| hsa-miR-7-5p | 95 | 0,00092033 | 0,906862745 | EPB41L2;ZBTB38;EHD3;FMNL3 TRIM44;ANKRD33B;IGFBP4 CLMP;FNBP1;PDCD1LG2 | 3,05516 | 0,01234 |
| hsa-miR-342-3p | 23 | 0,010804768 | 0,781862745 | DBN1;CCDC88A;SMARCA4;WDR3 CBX2;N4BP3;DNMBP;PWP2; MEX3A;PITPNM2 | -4,13469 | 0,037616 |
| hsa-miR-150-5p | 14 | 0,003273093 | 0,870098039 | SALL2;MEX3A;NARS2; KLHL7;UBE3D;ZNF11; TFAP4;FOXRED2;PDZD2;C17orf75 | -11,2608 | 0,016026 |
| hsa-miR-520d-5p+hsa-miR-527+hsa-miR-518a-5p | 4 | 0,023488652 | 0,713235294 | ZNF367;DNMT3B; ANKRD50;SPIN4 | -2,43353 | 0,0446 |
| hsa-miR-4443 | 2 | 0,033010244 | 0,676470588 | TBL2;SLC25A37 | -2,04241 | 0,034973 |
| hsa-miR-450b-5p | 2 | 0,043800186 | 0,637254902 | SLC25A37;SOX12 | -8,07801 | 0,037616 |
| hsa-miR-133b | 1 | 0,018633156 | 0,735294118 | KCNH2 | -2,6926 | 0,037616 |

#### 2. BuAL versus BuLY

The top 10 inverse-significant miRNA–mRNA interactions ranked by the number of validated DE targets observed among inverse-significant correlations, with correlation and evidence statistics retained as supporting metadata in BuAL versus BuLY are shown in table 2.

**Table 2.**
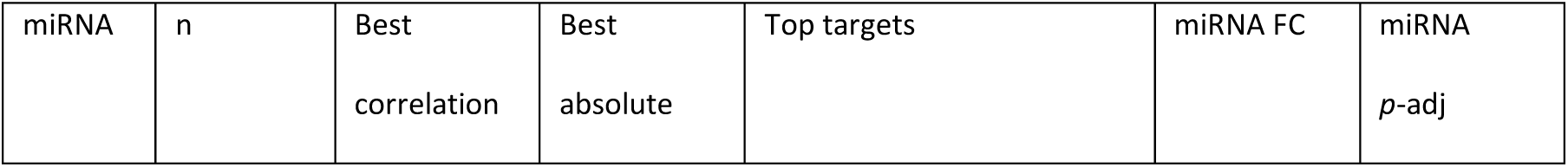
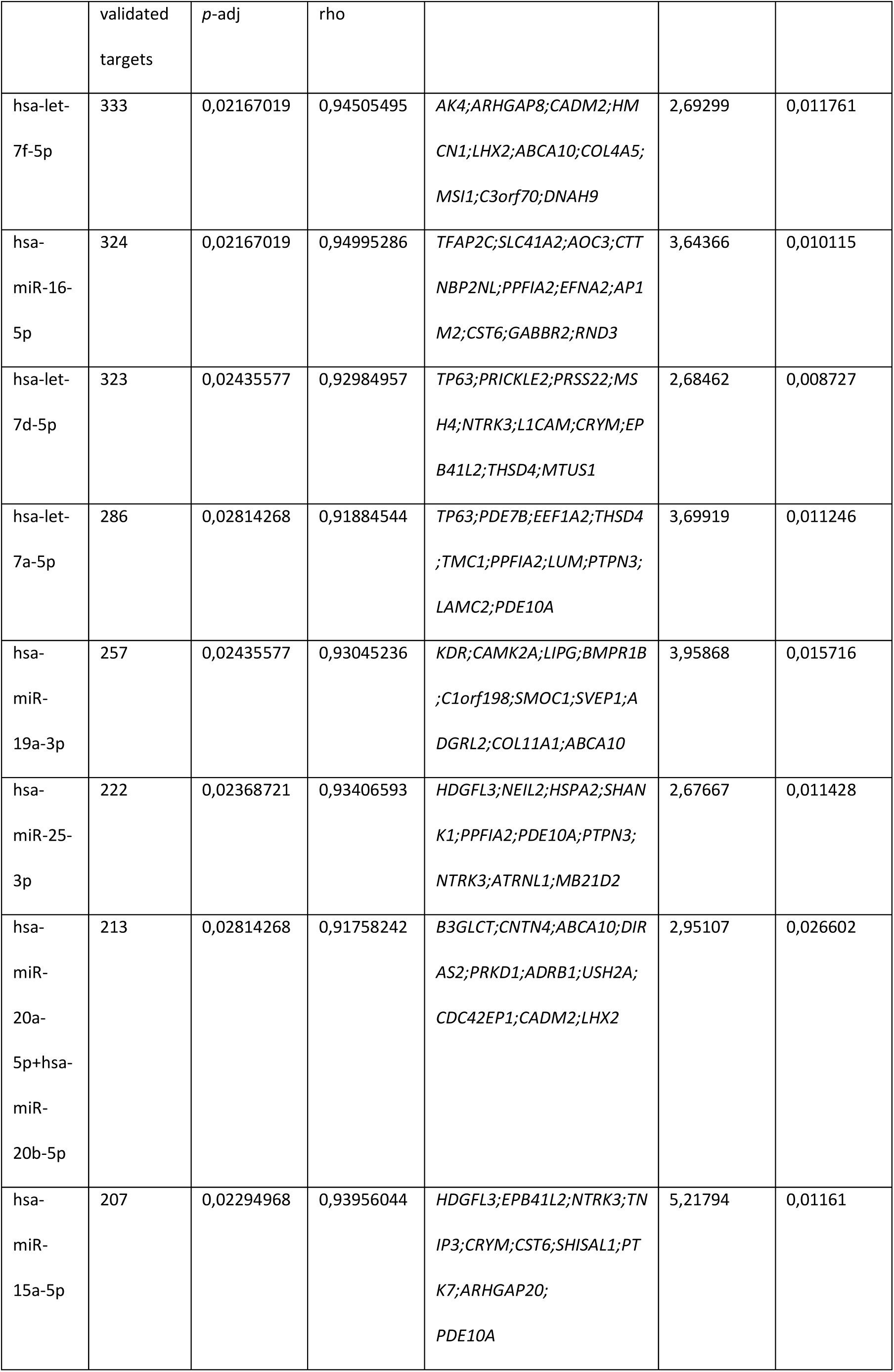
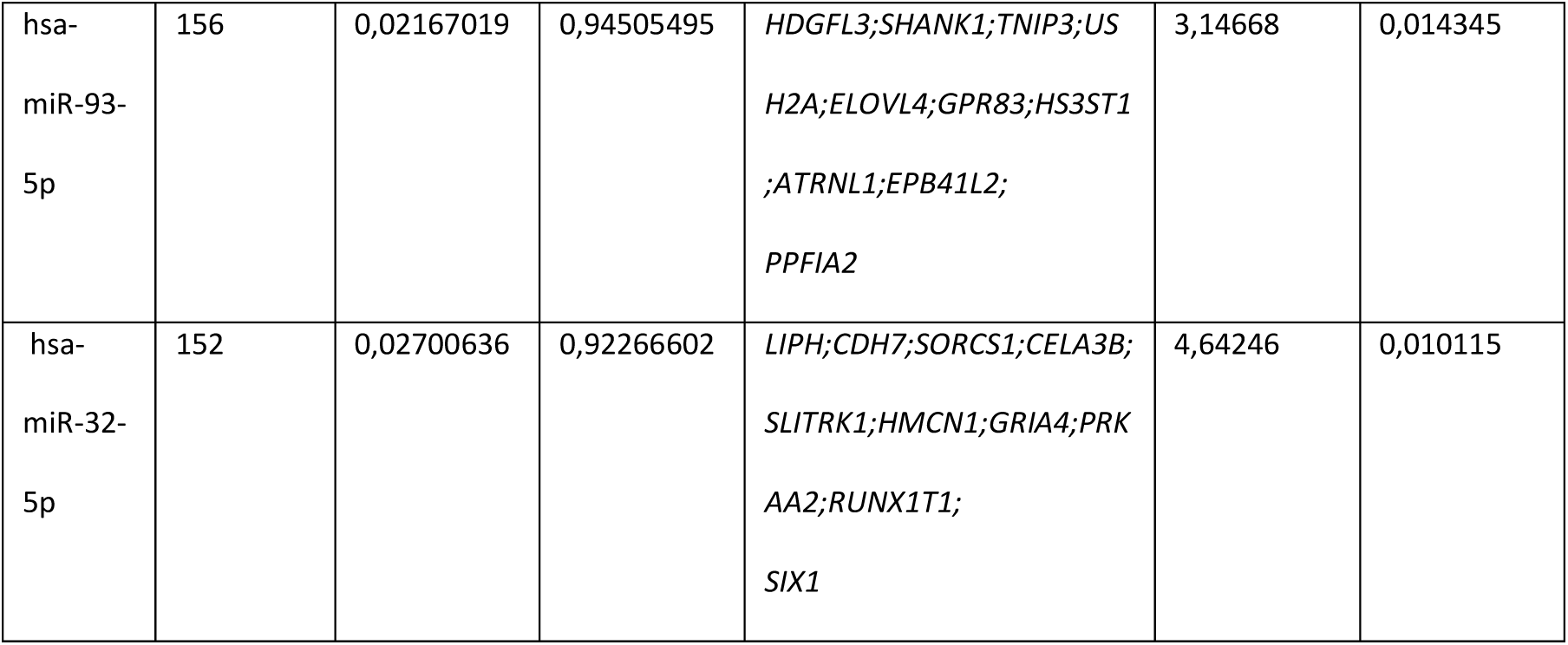
Top 10 Inverse-significant miRNA-mRNA interactions ranked by number of validated DE targets observed in BuAL versus BuLY.

All ranked inverse-significant miRNA-mRNA interactions for the BuAL vs BuLY comparison are shown in supplementary file 7. Edge-level validated, inverse-significant miRNA–mRNA interactions for the BuAL versus BuLY comparison (supplementary file 8) visualized as a MiRNA target network (STRING version 12.0) are shown in Figure 6a. Functional enrichment analysis of a total of 4085 inversely-expressed validated target genes of the DE miRNAs in BuAL compared to BuLY provided us with an overall clue of their functional roles (supplementary files 9 and 10). Whereas the 849 upregulated edge-level validated targets of DE downregulated miRNAs in BuAL versus BuLY (hsa-miR-34a-5p, hsa-let-7e-5p, hsa-miR-182-5p, hsa-miR-101-3p, hsa-miR-103a-3p, among others) are significantly associated with innate immune response pathways (Figure 6b), the 3286 downregulated edge-level validated targets of DE upregulated miRNAs in BuAL versus BuLY (hsa-let-7f-5p, hsa-miR-16-5p, hsa-let-7d-5p, hsa-let-7a-5p, hsa-miR-19a-3p, hsa-miR-25-3p, hsa-let-7g-5p, among others) are significantly related to cell adhesion, extracellular matrix organization and axon guidance pathways (FDR<0.05) (Figure 6c).

**Figure 6.**
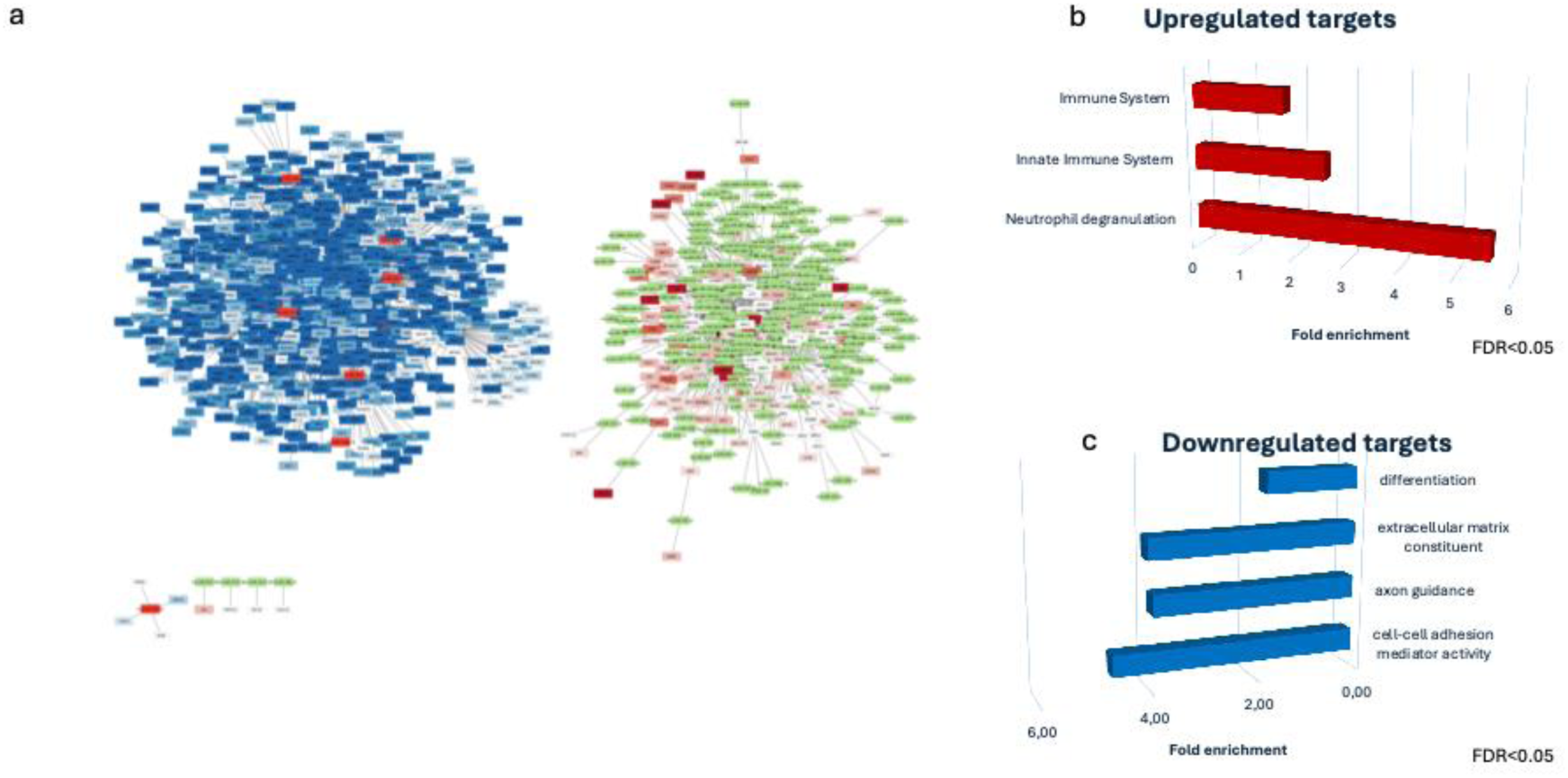
a_miRNA-mRNA target network analysis in BuAL versus BuLY comparison performed with STRING (version 12.0) including DEmiRNAs (FC > 1.5; p-adj < 0.05) and their validated targets according to the multiMiR R package; log_2_FC>|0.5|; p-adj<0.05). Only significant inverse associations were retained (ρ≤−0.5 and BH-adjusted correlation p-value<0.05). Upregulated miRNAs in red color, downregulated miRNAs in green color. Upregulated validated mRNA targets with significant edges in red color, downregulated validated mRNA targets with significant edges blue color. **b**_**DAVID Functional enrichment analysis of upregulated targets** (red color tones) fulfilling edge-level validated, inverse-significant miRNA–mRNA interactions (FDR<0.05). **c**_**DAVID Functional enrichment analysis of downregulated targets** (blue color tones) fulfilling edge-level validated, inverse-significant miRNA– mRNA interactions (FDR<0.05).

## Discussion

This deep mRNA sequencing analysis and integrative miRNA-mRNA network analysis provides further insight into the transcriptome and the molecular programs driving pediatric Burkitt leukemia and lymphoma. The transcriptomic differences observed between BL and normal lymphoid tissue inferred by functional enrichment analysis underscore a MYC-driven backbone with strong E2F4 and MYB pathway activity, activation of the MAPK and PI3K signaling pathways and repression of the JAK–STAT, NFkB, TP53, TRAIL and Hypoxia pathways, suggesting a strong proliferative core axis with attenuated cytokine, apoptotic, and stress-response signaling relative to normal lymphoid tissue. Consistent with the observation by other groups, in addition to deregulated MYC signaling, activation of the MAPK pathway(43), PI3K pathway(15, 44–46) and the MYC-miR-150-MYB pathway are key signaling pathways that contribute to BL pathogenesis(47). Loss of *TP53* promotes *MYC*-induced transformation by blocking the apoptosis that results when MYC oncoprotein is expressed in primary B cells(48). Recent studies have shown that *TP53* mutations have high prognostic value in BL(14, 49), particularly in conjunction with gains of GPC5/MIR17HG(12).

A further finding of this study is an increased metabolic activity in BL characterized by enriched oxidative phosphorylation, which has been involved in chemotherapy resistance in various other lymphoproliferative diseases(50, 51).

Additionally, the suppression of cytokine-mediated immune communication within the tumor microenvironment and attenuated inflammatory signaling observed in BL compared to normal lymphoid tissue strengthens the notion that the tumor itself harbors gene signatures that enable evasion of immune recognition(52).

In the present study miRNA signatures of relevance in BL compared to normal lymphoid tissue were identified and were evaluated for validated inverse-significant miRNA–mRNA correlations. The miRNA-mRNA integrative analysis led to further insight into a post-transcriptional fine-tuning of distinct proliferative, immune, and structural modules through DE miRNAs in BL compared to normal lymphoid tissue, showing that DE downregulated miRNAs in BL, including hsa-miR-342-3p and hsa-miR-150-5p, among others, inversely correlate with DE upregulated mRNA targets functionally related to activation of transcriptional activity and nuclear organization pathways, whereas DE upregulated miRNAs, including hsa-miR-106a-5p, hsa-miR-17-5p and hsa-miR-7-5p, inversely correlate with DE mRNA targets related to downregulated immune-response and T-cell activation in BL.

Differences between BuAL and BuLY subtypes have also been reported by other groups at the genomic(12) and transcriptomic(53) level. In this study, when comparing BuAL subtype to BuLY, an enrichment in innate defense pathways with heightened inflammatory, angiogenic, and cytokine-related activity in the leukemic phenotype as well as downregulation of genes involved in extracellular matrix organization, cell-adhesion and axon guidance pathways was observed, indicating a loss of adhesion and microenvironmental structure in the leukemic subtype relative to BuLY. Notably, the cascades with concordant behavior across both BuAL and BuLY subtypes include consistent activation of MYC, E2F, oxidative phosphorylation, reactive O2 species as well as MAPK signaling, sustaining a proliferative backbone, and downregulated TP53 and TRAIL pathways in both, indicating shared attenuation of apoptosis-related responses across the leukemic and lymphomatous subtypes in BL. Limitations to this study include a low patient number, limiting the robustness of the results of this pilot study, which is comparable in number of cases to other miRNA-mRNA interaction studies. Additionally, evaluation through miRNA sequencing may provide insight into further undetected miRNAs not covered by the miRNA microarray used. Nevertheless, in the present study a comprehensive analysis through mRNA-deep sequencing and miRNA-microarrays allowed us to identify transcriptional pathways and miRNA-mRNA networks of relevance in BL.

## Conclusions

In summary, this transcriptional pathway activity analysis provides new insights on key proliferative, metabolic and cytokine-mediated immune signaling pathways in BL as well as distinct miRNA signatures and miRNA-mRNA networks that modulate transcriptional activity and immune surveillance in BL. The results of this exploratory kick-off study shall pave the way for evaluation in larger cohorts and for functional studies.

## Declarations

### Ethics approval

Ethics Committee of the University of Rostock, Germany (Reference number A 2023-0054) and Ethics Committee of the Hospital Sant Joan de Déu, Barcelona, Spain (Reference number HSJD C.I. PIC-36-23).

Consent for publication: not applicable

### Availability of data and materials

Data are provided within the manuscript and supplementary information files. The datasets generated and analyzed during the current study are available in the NCBI’s Gene Expression Omnibus, are accessible through the GEO series accession numbers GSE337108 and GSE 337806 upon acceptance and through the corresponding author on request.

### Authors’ contribution statement

TC-S, VC, BS and CFC designed the study, CFC and TC-S acquired funding, TC-S, CR and BS performed the research, E-I V and TC-S performed bioinformatic and statistical analysis, VC, OS, EE-C, SM, SP, NC, MC, CM, CJ, JM and AMB contributed with study material and data. TC-S and E-I V wrote the article. All authors have read and approved the manuscript.

## List of abbreviations

BL: Burkitt leukemia and lymphoma
mRNA: messenger ribonucleic acid
miRNA: micro ribonucleic acid
BuAL: Burkitt leukemia
BuLY: Burkitt lymphoma
NCLT: normal control lymphoid tissue
DE: differential expression
DEG: differentially expressed genes
Hsa: homo sapiens
BH: Benjamini-Hochberg
NHL: non-Hodgkin Lymphoma
GCB: Germinal center B
BM: bone marrow
UTR: untranslated region
NCLT: normal control lymphoid tissue
EBER: Epstein-Barr virus-encoded small RNA
FDR: false discovery rate
FC: fold change
GO: gene ontology
KEGG: Kyoto Encyclopedia of Genes and Genomes
ORA: over-representation analyses
GSEA: gene set enrichment analysis
NES: normalized enrichment scores
TF: transcription factor
GO-BP: GO biological process

## Acknowledgements

We are indebted to the “Biobanc de lHospital Infantil Sant Joan de Déu per a la investigació” integrated in the Spanish Biobank Network of ISCIII for the sample and data procurement. We would like to acknowledge Novogene® for mRNA-seq support and Genevia Technologies for their help in statistical analysis and in constructing the figures and tables.

