## Supplementary methods and supplementary figures for "“mRNA deep sequencing profiling and integrative miRNA-mRNA network analysis in pediatric Burkitt leukemia and lymphoma “"

**mRNA-sequencing:**

Library preparation for Transcriptome sequencing:

Non strand specific library: mRNA was purified from total RNA using poly-T oligo-attached magnetic beads. After fragmentation, the first strand cDNA was synthesized using random hexamer primers followed by the second strand cDNA synthesis. The library was ready after end repair, A-tailing, adapter ligation, size selection, amplification, and purification. The library was checked with Qubit and real-time PCR for quantification and bioanalyzer for size distribution detection.

Clustering and sequencing: After library quality control, different libraries were pooled based on the effective concentration and targeted data amount, then subjected to Illumina sequencing.

Bioinformatics Analysis:

Data quality control: Raw data (raw reads) of fastq format were firstly processed through fastp software. Clean data (clean reads) were obtained by removing reads containing adapter, reads containing poly-N and low quality reads from raw data. Q20, Q30 and GC content of the clean data were calculated. A total of 19 samples were analyzed in this sequencing for transcriptome analysis, resulting in 377.46 Gb of clean bases. Each sample produced over 8.73G of clean bases, with the Q20 base percentage greater than 95.45 and the Q30 base percentage greater than 89.66 for each sample. All the downstream analyses were based on the clean data with high quality. Data can be accessed at the GSE337108 accession number.

Reads mapping to the reference genome:

HISAT2 (2.2.1) was used for mapping of reads to the reference genome.

Quantification of gene expression level: Feature Counts v1.5.0-p3 was used to count the reads numbers mapped to each gene. Fragments Per Kilobase of transcript sequence per Millions base pairs sequenced (FPKM) of each gene was calculated based on the length of the gene and reads count mapped to this gene.

Exploratory data assessment:

Exploratory data assessment included evaluation of sequencing depth, inter-sample correlations, and unsupervised variance structure. Total library size per sample was derived from raw counts and visualized using customized barplots and the ggplot2 R package (4.0.0), with cohort median and heuristic low-depth thresholds (10 million reads or median of 1.5 × interquartile range (IQR)) indicated for reference. Library sizes ranged from approximately 10 million to more than approximately 200 million reads, with all samples surpassing minimal inclusion criteria (Supplementary Figure 1). Differential expression modeling incorporated DESeq2 size-factor normalization to account for sequencing depth and compositional differences across libraries. For exploratory visualization and downstream correlation-based summaries, variance-stabilized expression values (VST) were used to reduce heteroscedasticity and improve comparability across samples. While lower-depth libraries may reduce sensitivity for lowly expressed transcripts, all samples passed basic QC thresholds and were retained.

Next, principal component analysis (PCA) was conducted on variance-stabilized expression values, restricted to the 500 most variable protein-coding genes (Supplementary Figure 2), while sample-to-sample Pearson correlations were summarized as a clustered heatmap (Supplementary Figure 3), using the pheatmap R package (1.0.13). To systematically identify potential outliers, a robust PCA was additionally performed using the PcaHubert algorithm from the rrcov (1.7-7), applying the 99th percentile of orthogonal and chi-square–derived score distances as detection thresholds (Supplementary Figure 4). Collectively, two pleural-fluid–derived Burkitt lymphoma samples (TC38, TC40) displayed modestly lower sequencing depth and slightly elevated variance metrics but remained within acceptable limits and did not exert disproportionate leverage. Both were therefore retained for all downstream analyses as biologically representative, lower-depth replicates.

**miRNA profiling:**

<u>miRNA arrays:</u>

Analysis of miRNA expression was performed using the NanoString nCounter System with the Human v3 miRNA assay (Nanostring, Seattle, WA, USA), analyzing 800 miRNAs per sample following the manufacturer´s instructions. 100 ng of total RNA was included per sample.

Raw data were imported from the nCounter Digital Analyzer and analyzed with the Rosalind® software (https://rosalind.bio/) using standard settings for background subtraction and normalization. Statistical analysis: Following background subtraction based on probe correction factors, normalization was performed in two steps, including positive control normalization and codeset normalization to adjust for batch effects. Group-wise comparisons with the t-test method were performed, delivering fold changes and p-values. P-value adjustment was performed through the Benjamini-Hochberg method of estimating false discovery rates (FDR). Clustering of miRNAs for the final heatmap of differentially expressed miRNA was done using the PAM (Partitioning Around Medoids) method.

miRNA validation

Four miRNAs among the most significantly up-and downregulated in BuAL compared to BuLY were chosen for exemplarily validation of miRNA array screening results by qPCR using miRNA-specific TaqMan Assays (Applied Biosystems, Darmstadt, Germany), containing RT-primers specific for the mature miRNAs. RNU6B was used as endogenous control (Applied Biosystems) in the entire cohort. The reverse transcription was carried out as a multiplex reaction, containing the RT-primers for all four miRNAs and the endogenous control in one reaction. The subsequent qPCR reactions were set up according to the manufacturer's protocol using TaqMan Universal PCR Master Mix II, No UNG (Applied Biosystems) and the miRNA-specific primer/probe mixes in singleplex reactions. The runs were performed in triplicates on a StepOne Plus Real-time PCR system (Applied Biosystems) and the data analyzed using the StepOne Software v2.3 (Applied Biosystems). Relative expression against RNU6B (fold change) was calculated using the $2^{-\Delta Ct}$-algorithm (supplementary table 3, supplementary Figure 5). Statistical significance analysis of comparisons between groups was assessed through the U-Mann-Whitney test.

**<u>Cell lines:</u>** DG-75 (Leibniz Institute DSMZ, Braunschweig, Germany) and Raji cells (ATCC, Manassas, VA, USA), authentication validated, were maintained in RPMI 1640 (PAN Biotech, Aidenbach, Germany) supplemented with 10% heat-inactivated fetal calf serum (Biochrom, Berlin, Germany) and 100 µg/mL penicillin and streptomycin (Biochrom, Berlin, Germany) at 37°C in a humidified 5% CO2-incubator. Mycoplasma contamination was excluded through periodic screening.

**Supplementary Figures**

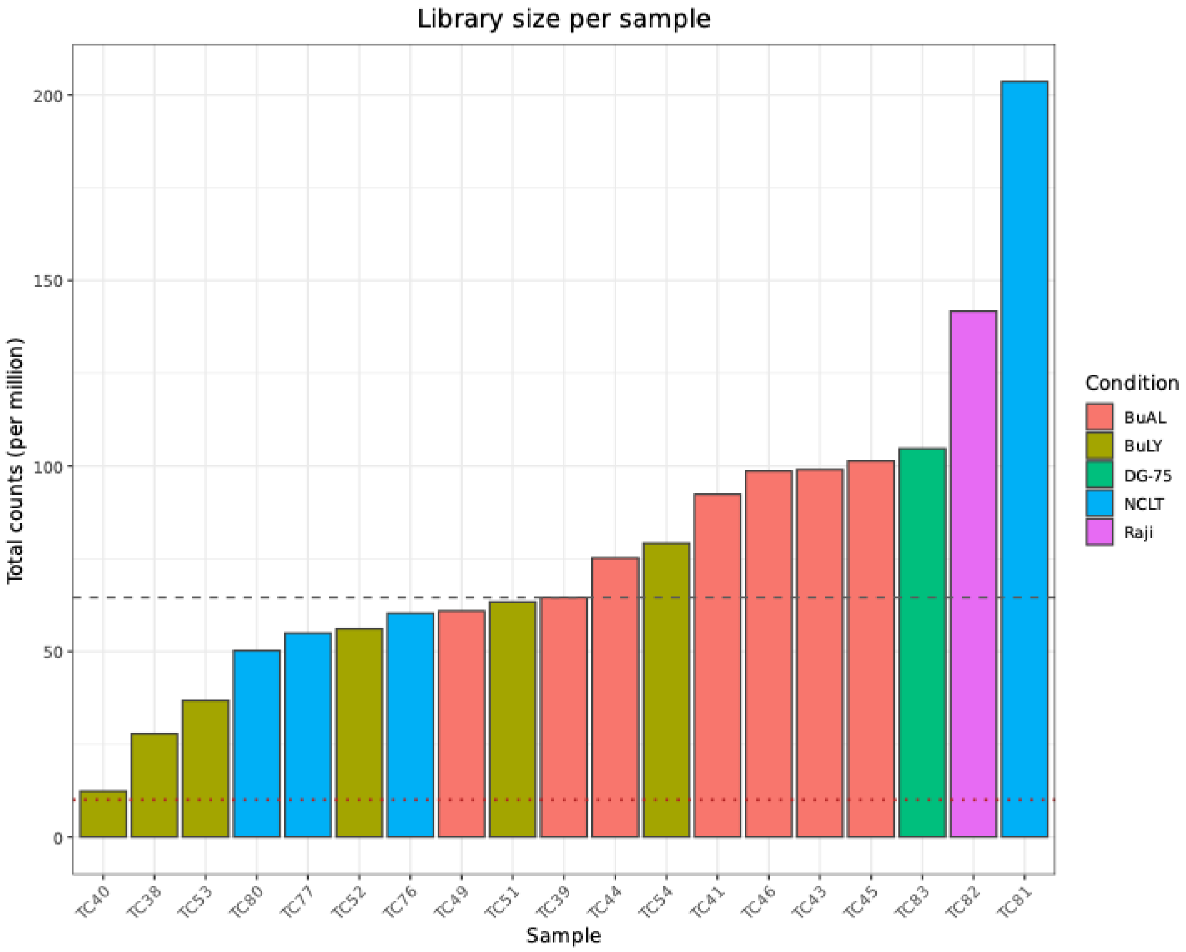


**Supplementary Figure 1: Barplot of total library size per RNA-seq sample**. Customized barplot generated with the ggplot2 R package (4.0.0) showing the total number of mapped reads per sample (in millions). Bars are colored by the main biological condition. The dashed line indicates the cohort median library size, while the dotted line marks an approximate low-depth threshold (set at the greater of 10 million reads or the median minus 1.5 × IQR). Samples falling near or below this reference are considered lower-depth libraries but remain part of the dataset unless additional QC metrics suggest otherwise.

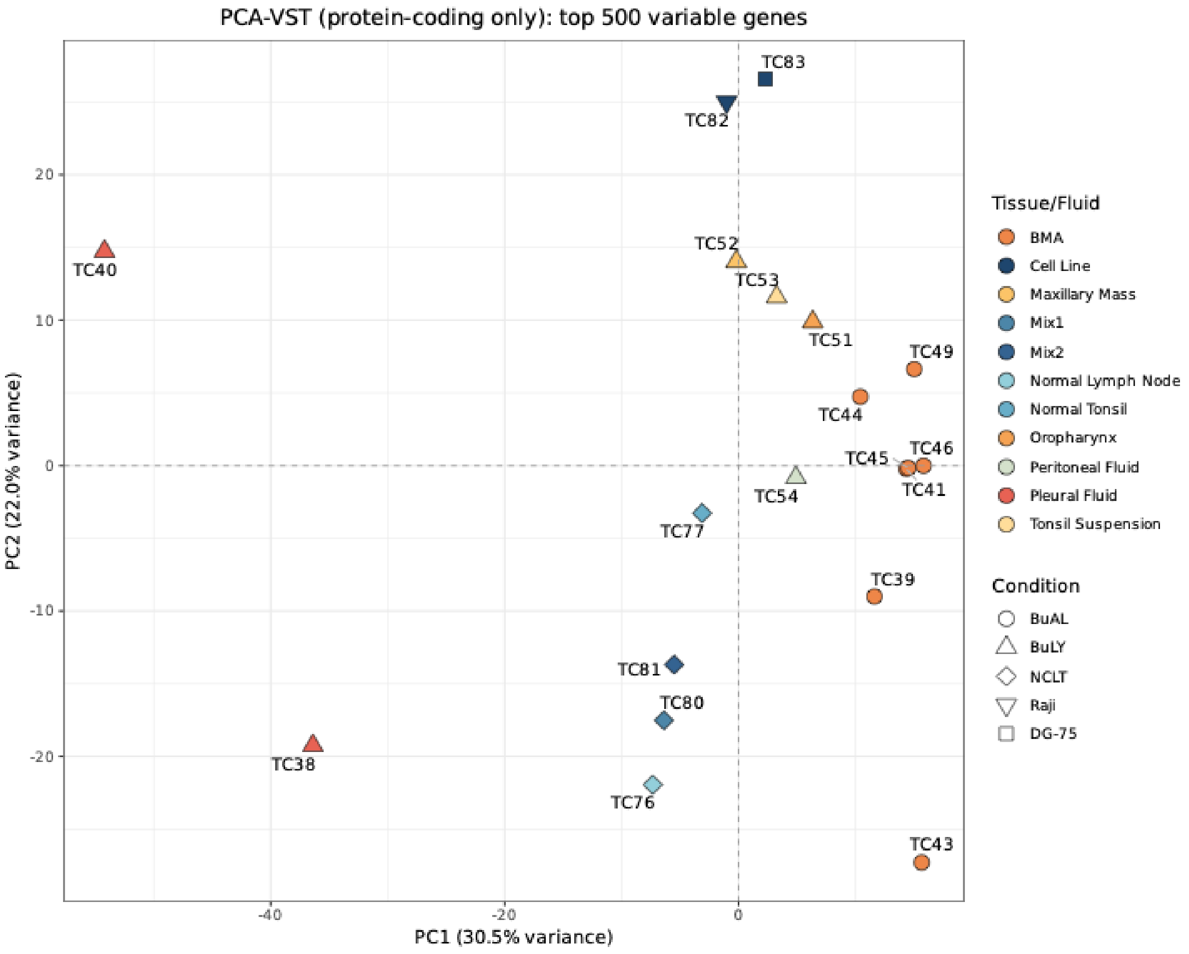


**Supplementary Figure 2: Principal component analysis (PCA) of RNA-seq samples.**
Principal component analysis (PCA) based on variance-stabilized (VST) expression values from the top 500 most variable protein-coding genes. Each point represents one sample, colored by biological condition. Tumor (BuAL, BuLY) and control (NCLT) samples separate along PC1 (30.5 % variance), indicating strong global expression differences between malignant and normal contexts. Two pleural-fluid–derived BuLY samples (TC38, TC40) showed higher dispersion along PC1 but remained within acceptable variance limits.

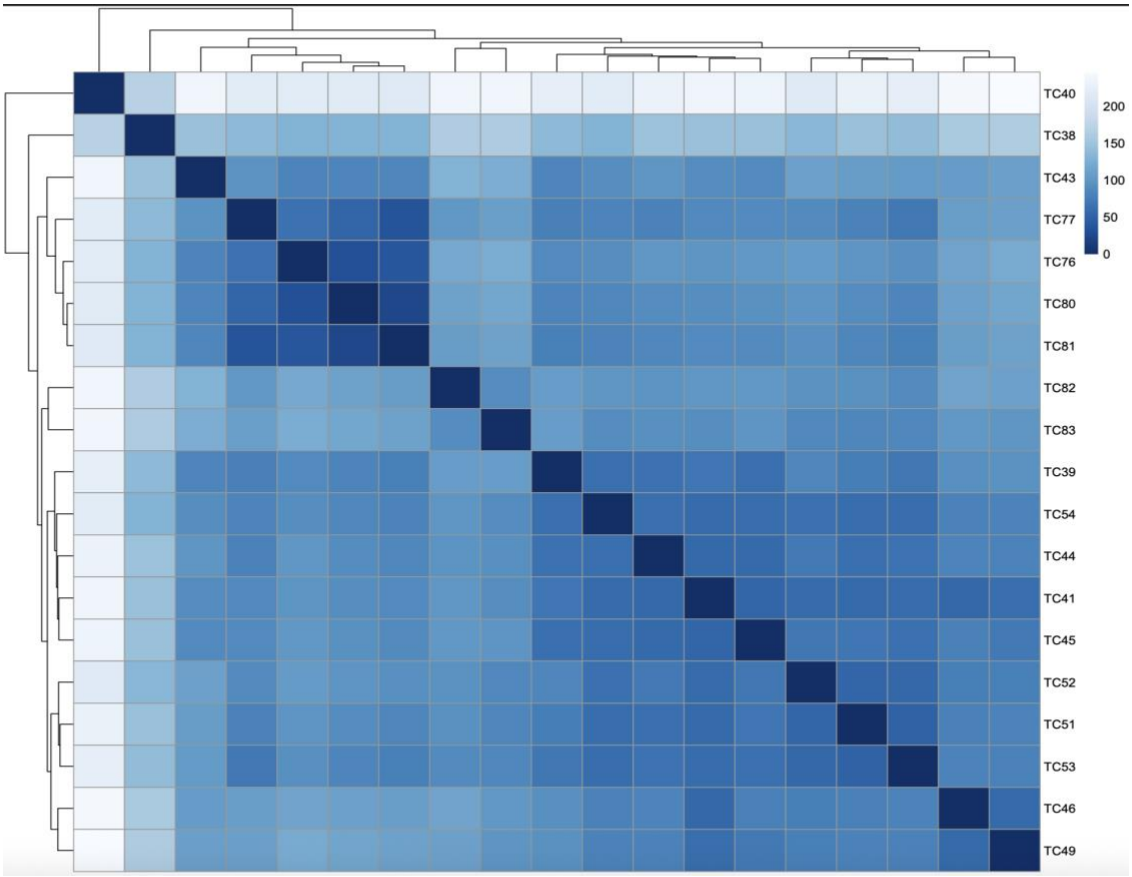


**Supplementary Figure 3: Sample-to-sample correlation heatmap.**
Hierarchical clustering heatmap of pairwise Pearson correlations computed from variance-stabilized expression values (protein-coding genes only), using the pheatmap R package (1.0.13).

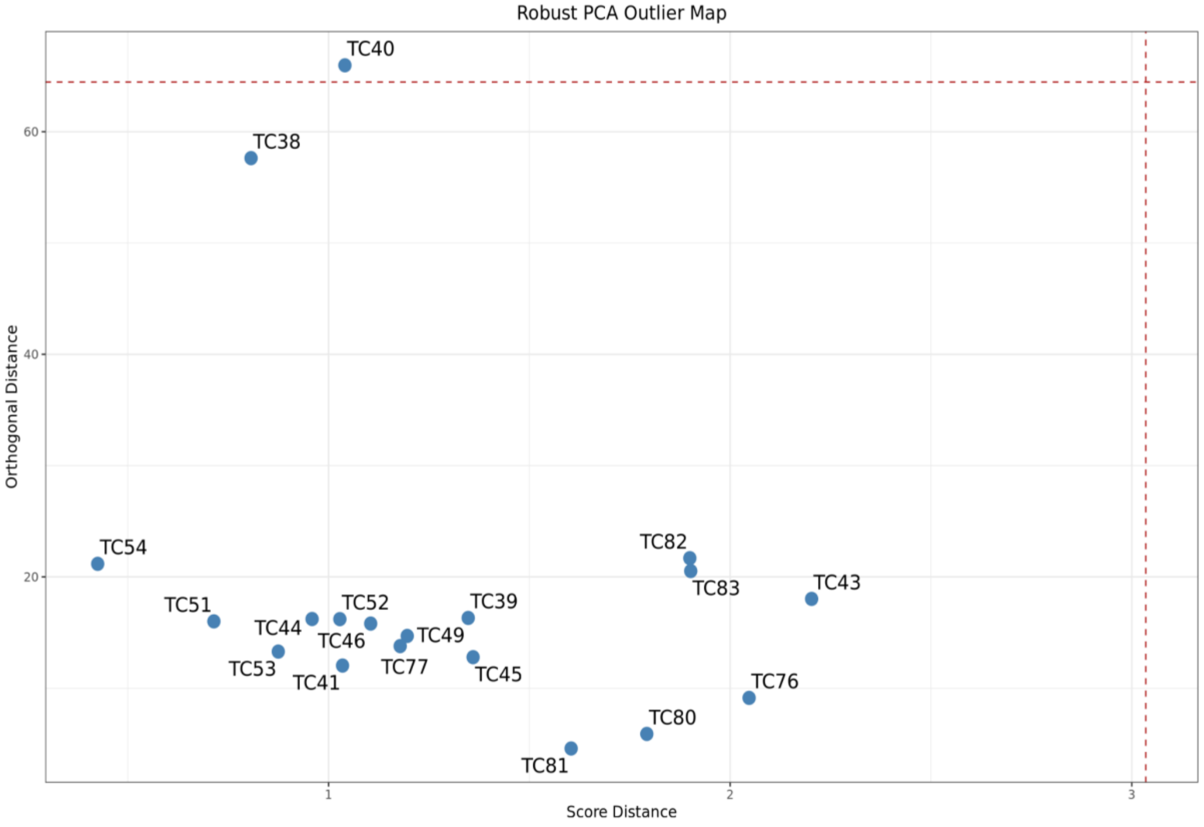


**Supplementary Figure 4: Robust PCA–based outlier detection across RNA-seq samples.** Outlier assessment based on a robust principal component analysis (PcaHubert) applied to variance-stabilized expression values of the 500 most variable protein-coding genes, using the rrcov R package (1.7-7). Each point represents one sample, positioned according to its score distance (x-axis) and orthogonal distance (y-axis). Dashed red lines denote the 99th-percentile cutoffs used to define potential outliers. Sample TC40 exceeded the orthogonal distance threshold, while TC38 was near but within limits; both samples were retained following manual review and depth verification.

a)

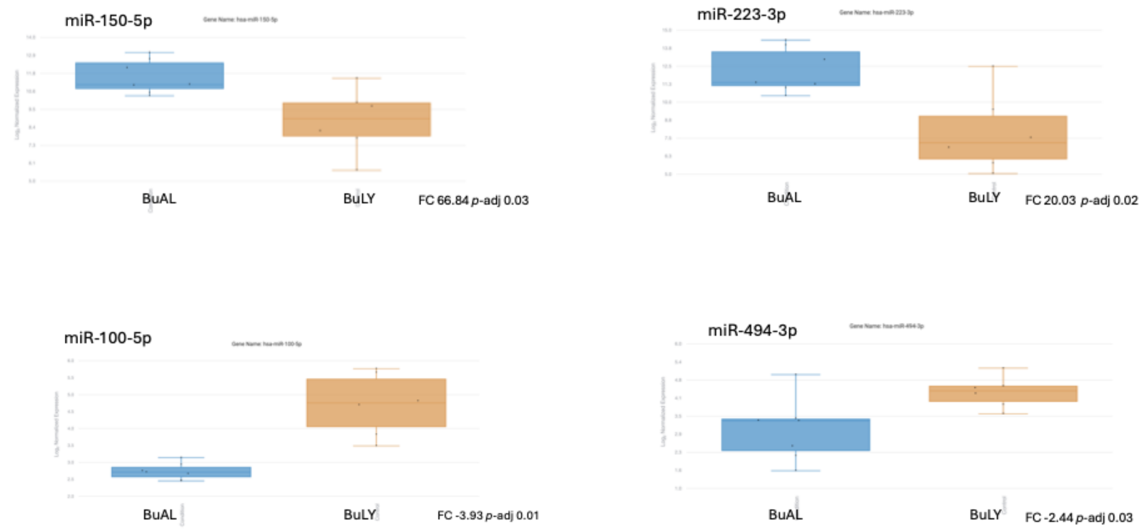


b)

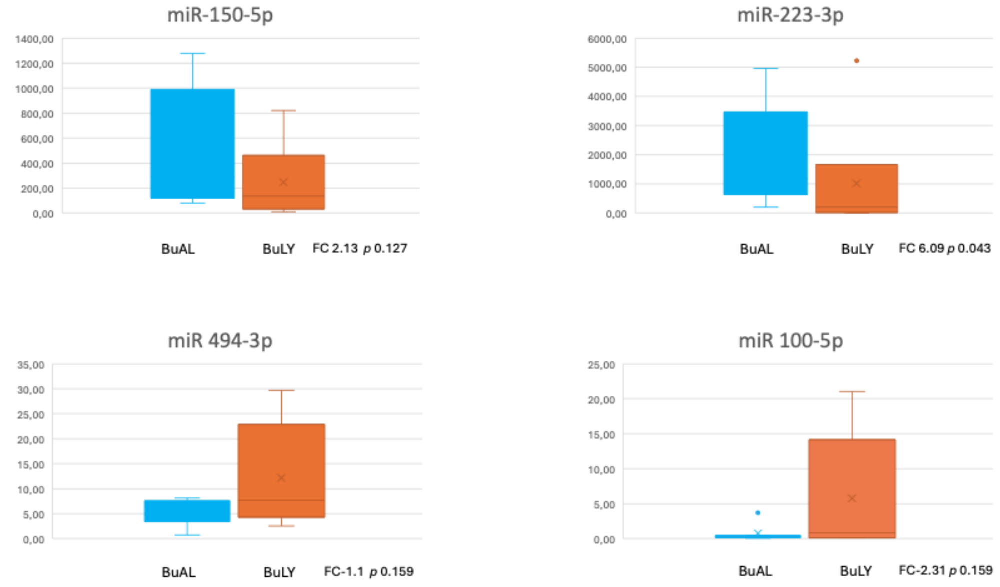


**Supplementary Figure 5: miRNA validation by qPCR analysis**

(a) Box-plots. Log2 normalized expression of miR-150-5p, miR-223-3p, miR-494-3p and miR-100-5p in Nanostring miRNA microarrays in BuAL versus BuLY. (b) qPCR analysis. miR-223-3p, miR-150-5p, miR-100-5p and miR-494-3p relative expression to RNU6B in BuAL versus BuLY.
