## Supplementary tables for "“mRNA deep sequencing profiling and integrative miRNA-mRNA network analysis in pediatric Burkitt leukemia and lymphoma “"

Supplementary tables 1-3

Supplementary table 1 miRNAs differentially expressed in BL (n=13) versus NCLT (n=4) (FC>1.5, *p*-adj<0.05)

Supplementary table 2 Differentially expressed miRNAs in BuAL (n=7) and BuLY (n=6) (FC>1.5, *p*-adj<0.05)

Supplementary table 3 qPCR analysis for miR-223-3p, miR-150-5p, miR-100-5p and miR-494-3p against the endogenous control RNU6B

Supplementary table 1_ miRNAs differentially expressed in BL (n=13) versus NCLT (n=4). (FC>1.5, *p*-adj<0.05)

| B-AL/B-Ly (n=13) versus NCLT (n=4)<br>Name | Fold Change | p-Value | p-Adj |
|---|---|---|---|
| hsa-miR-106a-5p+hsa-miR-17-5p | 4.73338 | 5.72e-05 | 0.016026 |
| hsa-miR-7-5p | 3.05516 | 1.55e-05 | 0.012340 |
| hsa-miR-150-5p | -11.2608 | 6.02e-05 | 0.016026 |
| hsa-miR-450b-5p | -8.07801 | 0.000330 | 0.037616 |
| hsa-miR-342-3p | -4.13469 | 0.000313 | 0.037616 |
| hsa-miR-133b | -2.6926 | 0.000278 | 0.037616 |
| hsa-miR-520d-5p+hsa-miR-527+hsa-miR-518a-5p | -2.43353 | 0.000447 | 0.044600 |
| hsa-miR-4443 | -2.04241 | 0.000175 | 0.034973 |

Supplementary table 2_ Differentially expressed miRNAs in BuAL (n=7) and BuLY (n=6) (FC>1.5, *p*-adj<0.05)

| Name | Fold Change | p-Value | p-Adj |
|---|---|---|---|
| hsa-miR-223-3p | 20.0324 | 0.009643 | 0.017189 |
| hsa-miR-144-3p | 17.7322 | 0.009984 | 0.009984 |
| hsa-miR-142-3p | 9.71464 | 0.002396 | 0.010150 |
| hsa-miR-150-5p | 6.8366 | 0.018472 | 0.026753 |
| hsa-miR-15a-5p | 5.21794 | 0.003893 | 0.011610 |
| hsa-miR-32-5p | 4.64246 | 0.002095 | 0.010115 |
| hsa-miR-29b-3p | 4.02188 | 0.005190 | 0.012837 |
| hsa-miR-19a-3p | 3.95868 | 0.008271 | 0.015716 |
| hsa-miR-374a-5p | 3.7809 | 0.002669 | 0.010150 |
| hsa-let-7a-5p | 3.69919 | 0.003523 | 0.011246 |
| hsa-miR-16-5p | 3.64366 | 0.002106 | 0.010115 |
| hsa-let-7g-5p | 3.59163 | 0.000125 | 0.008727 |
| hsa-miR-15b-5p | 3.42115 | 0.011337 | 0.019255 |

| | | | |
|---|---|---|---|
| hsa-miR-19b-3p | 3.1736 | 0.026953 | 0.035728 |
| hsa-miR-93-5p | 3.14668 | 0.006705 | 0.014345 |
| hsa-miR-20a-5p+hsa-miR-20b-5p | 2.95107 | 0.018268 | 0.026602 |
| hsa-miR-98-5p | 2.8632 | 0.000620 | 0.009984 |
| hsa-miR-106a-5p+hsa-miR-17-5p | 2.78535 | 0.018223 | 0.026585 |
| hsa-let-7f-5p | 2.69299 | 0.003984 | 0.011761 |
| hsa-let-7d-5p | 2.68462 | 8.69e-05 | 0.008727 |
| hsa-miR-25-3p | 2.67667 | 0.003745 | 0.011428 |
| hsa-miR-191-5p | 2.42768 | 0.006891 | 0.014509 |
| hsa-miR-148a-3p | 2.27991 | 0.007755 | 0.015426 |
| hsa-miR-7-5p | 1.90303 | 0.000839 | 0.009984 |
| hsa-miR-140-5p | 1.8127 | 0.006044 | 0.013562 |
| hsa-miR-148b-3p | 1.65901 | 0.013654 | 0.021576 |
| hsa-miR-30d-5p | 1.56118 | 0.008888 | 0.016419 |
| hsa-miR-26b-5p | 1.5532 | 0.015215 | 0.023260 |
| hsa-miR-1285-5p | -8.74465 | 3.20e-05 | 0.008727 |
| hsa-miR-1972 | -7.314 | 0.000119 | 0.008727 |
| hsa-miR-548aa+hsa-miR-548t-3p | -5.91308 | 0.002006 | 0.010115 |
| hsa-miR-548ah-5p | -5.51557 | 5.40e-05 | 0.008727 |
| hsa-miR-4787-3p | -5.01173 | 0.001163 | 0.009984 |
| hsa-miR-320e | -4.94538 | 0.027968 | 0.036768 |
| hsa-miR-875-3p | -4.78895 | 4.71e-05 | 0.008727 |
| hsa-miR-575 | -4.63851 | 0.012774 | 0.020634 |
| hsa-miR-369-5p | -4.50354 | 0.000285 | 0.009488 |
| hsa-miR-3614-3p | -4.50295 | 0.000694 | 0.009984 |
| hsa-miR-3179 | -4.43951 | 0.001935 | 0.010115 |
| hsa-miR-1277-3p | -4.41833 | 0.006318 | 0.013889 |
| hsa-miR-152-5p | -4.33725 | 0.000555 | 0.009984 |
| hsa-miR-1245b-3p | -4.32788 | 0.002017 | 0.010115 |
| hsa-miR-224-5p | -4.292 | 0.000254 | 0.009488 |
| hsa-miR-1290 | -4.289 | 0.000370 | 0.009488 |
| hsa-miR-1233-3p | -4.27303 | 0.000320 | 0.009488 |
| hsa-miR-450b-3p | -4.272 | 0.002716 | 0.010150 |
| hsa-miR-652-5p | -4.25214 | 0.001995 | 0.010115 |
| hsa-miR-1305 | -4.19709 | 0.000380 | 0.009488 |
| hsa-miR-208a-3p | -4.19631 | 0.000338 | 0.009488 |
| hsa-miR-4488 | -4.18051 | 0.001864 | 0.010115 |
| hsa-miR-1185-2-3p | -4.17889 | 0.000638 | 0.009984 |
| hsa-miR-2113 | -4.0765 | 0.001119 | 0.009984 |
| hsa-miR-508-5p | -4.03794 | 0.000916 | 0.009984 |
| hsa-miR-656-3p | -4.00141 | 0.014097 | 0.021954 |

| | | | |
|---|---|---|---|
| hsa-miR-483-5p | -3.97568 | 0.004326 | 0.012091 |
| hsa-miR-429 | -3.97342 | 0.001388 | 0.009984 |
| hsa-miR-100-5p | -3.9294 | 0.002803 | 0.010222 |
| hsa-miR-181a-2-3p | -3.9214 | 0.001323 | 0.009984 |
| hsa-miR-515-5p | -3.91114 | 0.003314 | 0.011041 |
| hsa-miR-325 | -3.91036 | 0.012622 | 0.020526 |
| hsa-miR-519c-3p | -3.89702 | 0.001889 | 0.010115 |
| hsa-miR-381-5p | -3.87286 | 0.001680 | 0.010115 |
| hsa-miR-518c-3p | -3.8635 | 0.001718 | 0.010115 |
| hsa-miR-300 | -3.86289 | 0.001413 | 0.009984 |
| hsa-miR-4455 | -3.86174 | 0.000128 | 0.008727 |
| hsa-miR-1286 | -3.82251 | 0.010152 | 0.017767 |
| hsa-miR-888-5p | -3.80987 | 0.000233 | 0.009488 |
| hsa-miR-4707-5p | -3.79421 | 0.001099 | 0.009984 |
| hsa-miR-497-5p | -3.78806 | 0.001960 | 0.010115 |
| hsa-miR-604 | -3.78171 | 0.000443 | 0.009862 |
| hsa-miR-517c-3p+hsa-miR-519a-3p | -3.76863 | 0.003576 | 0.011294 |
| hsa-miR-301a-5p | -3.76325 | 0.002281 | 0.010148 |
| hsa-miR-3158-3p | -3.75924 | 0.002067 | 0.010115 |
| hsa-miR-219a-5p | -3.75067 | 0.000497 | 0.009984 |
| hsa-miR-3130-3p | -3.75044 | 0.006050 | 0.013562 |
| hsa-miR-216a-5p | -3.75031 | 0.015031 | 0.023023 |
| hsa-miR-346 | -3.74226 | 0.009743 | 0.017240 |
| hsa-miR-328-5p | -3.73133 | 0.000833 | 0.009984 |
| hsa-miR-933 | -3.72916 | 0.002212 | 0.010115 |
| hsa-miR-525-5p | -3.7209 | 0.002289 | 0.010148 |
| hsa-miR-576-3p | -3.71288 | 0.000164 | 0.008727 |
| hsa-miR-492 | -3.69717 | 0.003600 | 0.011309 |
| hsa-miR-409-5p | -3.69343 | 0.003168 | 0.010832 |
| hsa-miR-518e-3p | -3.67081 | 0.003752 | 0.011428 |
| hsa-miR-574-5p | -3.66443 | 9.24e-05 | 0.008727 |
| hsa-miR-3147 | -3.66352 | 0.003544 | 0.011267 |
| hsa-miR-1249-3p | -3.65938 | 0.002361 | 0.010150 |
| hsa-miR-593-3p | -3.65786 | 0.001590 | 0.010115 |
| hsa-miR-3182 | -3.65609 | 0.001812 | 0.010115 |
| hsa-miR-1288-3p | -3.65092 | 0.000738 | 0.009984 |
| hsa-miR-639 | -3.63526 | 0.006777 | 0.014388 |
| hsa-miR-548ar-5p | -3.61569 | 0.000105 | 0.008727 |
| hsa-miR-607 | -3.58947 | 0.000434 | 0.009862 |
| hsa-miR-34c-5p | -3.58447 | 0.004656 | 0.012469 |
| hsa-miR-567 | -3.57425 | 0.002143 | 0.010115 |

| | | | |
|---|---|---|---|
| hsa-miR-134-3p | -3.57336 | 0.002132 | 0.010115 |
| hsa-miR-5001-5p | -3.57088 | 0.000379 | 0.009488 |
| hsa-miR-30a-3p | -3.56425 | 0.002759 | 0.010194 |
| hsa-miR-4536-3p | -3.56381 | 0.001142 | 0.009984 |
| hsa-miR-2117 | -3.56245 | 0.000159 | 0.008727 |
| hsa-miR-599 | -3.55202 | 0.001847 | 0.010115 |
| hsa-miR-499a-3p | -3.55192 | 0.000599 | 0.009984 |
| hsa-miR-431-5p | -3.55126 | 0.012416 | 0.020344 |
| hsa-miR-510-3p | -3.54395 | 0.000812 | 0.009984 |
| hsa-miR-4284 | -3.53456 | 0.000733 | 0.009984 |
| hsa-miR-3168 | -3.53221 | 0.002440 | 0.010150 |
| hsa-miR-1297 | -3.53204 | 0.001643 | 0.010115 |
| hsa-miR-4787-5p | -3.53067 | 0.001120 | 0.009984 |
| hsa-miR-378d | -3.52991 | 0.004112 | 0.011888 |
| hsa-miR-1272 | -3.52983 | 0.002381 | 0.010150 |
| hsa-miR-205-5p | -3.52487 | 0.025669 | 0.034485 |
| hsa-miR-1185-5p | -3.52231 | 0.003059 | 0.010705 |
| hsa-miR-455-5p | -3.51672 | 0.001126 | 0.009984 |
| hsa-miR-182-3p | -3.51282 | 0.004708 | 0.012535 |
| hsa-miR-548v | -3.51134 | 0.002020 | 0.010115 |
| hsa-miR-153-3p | -3.50803 | 0.005027 | 0.012745 |
| hsa-miR-133a-3p | -3.50705 | 0.006890 | 0.014509 |
| hsa-miR-1205 | -3.49187 | 0.000163 | 0.008727 |
| hsa-miR-193a-3p | -3.4917 | 0.019567 | 0.027983 |
| hsa-miR-585-3p | -3.49156 | 0.003949 | 0.011714 |
| hsa-miR-520c-3p | -3.48948 | 0.002384 | 0.010150 |
| hsa-miR-187-3p | -3.48056 | 0.001809 | 0.010115 |
| hsa-miR-452-5p | -3.47948 | 0.004615 | 0.012443 |
| hsa-miR-541-3p | -3.47902 | 0.001614 | 0.010115 |
| hsa-miR-548y | -3.47733 | 0.001216 | 0.009984 |
| hsa-miR-937-3p | -3.47644 | 0.001174 | 0.009984 |
| hsa-miR-495-3p | -3.46763 | 0.001874 | 0.010115 |
| hsa-miR-648 | -3.46676 | 0.015604 | 0.023539 |
| hsa-miR-487b-5p | -3.46386 | 0.003382 | 0.011041 |
| hsa-miR-887-5p | -3.45724 | 0.005035 | 0.012745 |
| hsa-miR-514b-5p | -3.45187 | 0.001174 | 0.009984 |
| hsa-miR-550a-5p | -3.44953 | 0.000758 | 0.009984 |
| hsa-miR-339-3p | -3.41428 | 0.007895 | 0.015451 |
| hsa-miR-526b-5p | -3.41421 | 0.004248 | 0.012091 |
| hsa-miR-615-3p | -3.41179 | 0.001861 | 0.010115 |
| hsa-miR-6721-5p | -3.40811 | 0.011858 | 0.019789 |

| | | | |
|---|---|---|---|
| hsa-miR-200b-3p | -3.40527 | 0.002811 | 0.010222 |
| hsa-miR-448 | -3.40206 | 0.004773 | 0.012611 |
| hsa-miR-769-3p | -3.40051 | 0.004653 | 0.012469 |
| hsa-miR-210-5p | -3.39716 | 0.003447 | 0.011180 |
| hsa-miR-6720-3p | -3.39163 | 0.000555 | 0.009984 |
| hsa-miR-197-5p | -3.38722 | 8.37e-05 | 0.008727 |
| hsa-miR-654-3p | -3.38351 | 0.013819 | 0.021708 |
| hsa-miR-432-5p | -3.37613 | 0.002722 | 0.010150 |
| hsa-miR-6503-5p | -3.37101 | 0.002694 | 0.010150 |
| hsa-miR-4755-5p | -3.36919 | 0.004712 | 0.012535 |
| hsa-miR-1247-5p | -3.36692 | 0.000831 | 0.009984 |
| hsa-miR-616-3p | -3.36652 | 0.001814 | 0.010115 |
| hsa-miR-1200 | -3.36436 | 0.001319 | 0.009984 |
| hsa-miR-363-5p | -3.3624 | 0.003055 | 0.010705 |
| hsa-miR-1269a | -3.35795 | 0.005808 | 0.013205 |
| hsa-miR-1255a | -3.35788 | 0.002203 | 0.010115 |
| hsa-miR-566 | -3.35518 | 0.001964 | 0.010115 |
| hsa-miR-34a-5p | -3.34802 | 0.000784 | 0.009984 |
| hsa-miR-548e-5p | -3.32979 | 0.000417 | 0.009862 |
| hsa-miR-4485-3p | -3.32695 | 0.011879 | 0.019789 |
| hsa-miR-1295a | -3.32361 | 0.001034 | 0.009984 |
| hsa-miR-519b-5p+hsa-miR-519c-5p+hsa-miR-523-5p+hsa-miR-518e-5p+hsa-miR-522-5p+hsa-miR-519a-5p | -3.31457 | 0.005210 | 0.012837 |
| hsa-miR-488-3p | -3.31062 | 0.011825 | 0.019783 |
| hsa-miR-200a-3p | -3.30104 | 0.002262 | 0.010141 |
| hsa-miR-1908-3p | -3.29626 | 0.001436 | 0.009984 |
| hsa-miR-1262 | -3.28634 | 0.002231 | 0.010115 |
| hsa-miR-874-3p | -3.2836 | 0.004829 | 0.012635 |
| hsa-miR-1253 | -3.27944 | 0.001921 | 0.010115 |
| hsa-miR-487a-3p | -3.27919 | 0.003683 | 0.011354 |
| hsa-miR-208b-3p | -3.27892 | 0.003581 | 0.011294 |
| hsa-miR-513a-3p | -3.27503 | 0.003806 | 0.011504 |
| hsa-miR-521 | -3.27258 | 0.008230 | 0.015686 |
| hsa-miR-338-5p | -3.27195 | 0.005609 | 0.013097 |
| hsa-miR-524-3p | -3.26986 | 0.007915 | 0.015451 |
| hsa-miR-764 | -3.26499 | 0.005485 | 0.013033 |
| hsa-miR-3614-5p | -3.26459 | 0.001121 | 0.009984 |
| hsa-miR-4451 | -3.25765 | 0.010517 | 0.018285 |
| hsa-miR-124-3p | -3.24993 | 0.003330 | 0.011041 |
| hsa-miR-302a-5p | -3.2498 | 0.009765 | 0.017240 |
| hsa-miR-125b-5p | -3.24757 | 0.012511 | 0.020432 |

| | | | |
|---|---|---|---|
| hsa-miR-660-3p | -3.24514 | 0.006107 | 0.013650 |
| hsa-miR-487b-3p | -3.24284 | 0.000893 | 0.009984 |
| hsa-miR-873-3p | -3.24224 | 0.009040 | 0.016623 |
| hsa-miR-584-5p | -3.23869 | 0.004484 | 0.012321 |
| hsa-miR-1-3p | -3.23573 | 0.003099 | 0.010763 |
| hsa-miR-23c | -3.23436 | 0.002624 | 0.010150 |
| hsa-miR-626 | -3.23337 | 0.003169 | 0.010832 |
| hsa-miR-2682-5p | -3.23142 | 0.001750 | 0.010115 |
| hsa-miR-3065-3p | -3.22668 | 0.002346 | 0.010150 |
| hsa-miR-495-5p | -3.22295 | 0.004437 | 0.012268 |
| hsa-miR-369-3p | -3.22029 | 0.001411 | 0.009984 |
| hsa-miR-214-3p | -3.21638 | 0.011965 | 0.019892 |
| hsa-miR-211-5p | -3.2152 | 0.001718 | 0.010115 |
| hsa-mir-4532 | -3.2148 | 0.027005 | 0.035738 |
| hsa-miR-335-5p | -3.21393 | 0.001889 | 0.010115 |
| hsa-miR-196a-5p | -3.2124 | 5.25e-05 | 0.008727 |
| hsa-miR-548ar-3p | -3.20492 | 0.005951 | 0.013490 |
| hsa-miR-627-3p | -3.20372 | 0.001291 | 0.009984 |
| hsa-miR-149-5p | -3.20173 | 0.004137 | 0.011918 |
| hsa-miR-1296-3p | -3.19966 | 0.002457 | 0.010150 |
| hsa-miR-1197 | -3.1996 | 0.002908 | 0.010360 |
| hsa-mir-549a | -3.1957 | 0.003887 | 0.011610 |
| hsa-miR-939-5p | -3.19105 | 0.002423 | 0.010150 |
| hsa-miR-650 | -3.18647 | 0.002592 | 0.010150 |
| hsa-miR-4448 | -3.18618 | 0.000781 | 0.009984 |
| hsa-miR-1224-3p | -3.18415 | 0.002846 | 0.010222 |
| hsa-miR-548z+hsa-miR-548h-3p | -3.1778 | 0.000239 | 0.009488 |
| hsa-miR-601 | -3.17661 | 0.001779 | 0.010115 |
| hsa-miR-378f | -3.17265 | 0.002957 | 0.010443 |
| hsa-miR-33b-5p | -3.17109 | 0.004953 | 0.012745 |
| hsa-miR-556-5p | -3.17028 | 0.010510 | 0.018285 |
| hsa-miR-370-3p | -3.16102 | 0.001653 | 0.010115 |
| hsa-miR-184 | -3.1541 | 0.006629 | 0.014335 |
| hsa-miR-412-3p | -3.15242 | 0.000525 | 0.009984 |
| hsa-miR-513c-5p | -3.14895 | 0.017488 | 0.025844 |
| hsa-miR-133a-5p | -3.14554 | 0.008229 | 0.015686 |
| hsa-miR-301b-5p | -3.14524 | 0.001744 | 0.010115 |
| hsa-miR-499a-5p | -3.14382 | 0.010097 | 0.017709 |
| hsa-miR-135b-5p | -3.14337 | 0.020479 | 0.028976 |
| hsa-miR-181d-3p | -3.14293 | 0.003390 | 0.011041 |
| hsa-miR-27a-3p | -3.14206 | 0.001353 | 0.009984 |

| | | | |
|---|---|---|---|
| hsa-miR-1228-3p | -3.1354 | 0.003685 | 0.011354 |
| hsa-miR-642a-5p | -3.13284 | 0.001154 | 0.009984 |
| hsa-miR-548b-3p | -3.13251 | 0.011413 | 0.019255 |
| hsa-miR-513a-5p | -3.12813 | 0.010999 | 0.018795 |
| hsa-miR-876-5p | -3.12406 | 0.002656 | 0.010150 |
| hsa-miR-219a-1-3p | -3.12358 | 0.000799 | 0.009984 |
| hsa-miR-514a-3p | -3.11523 | 0.002250 | 0.010141 |
| hsa-miR-5001-3p | -3.10835 | 0.005655 | 0.013097 |
| hsa-miR-10a-5p | -3.10566 | 0.001641 | 0.010115 |
| hsa-miR-4647 | -3.09878 | 0.002212 | 0.010115 |
| hsa-miR-129-2-3p | -3.09835 | 0.007065 | 0.014708 |
| hsa-miR-517a-3p | -3.0962 | 0.002714 | 0.010150 |
| hsa-miR-3690 | -3.09616 | 0.004516 | 0.012321 |
| hsa-miR-25-5p | -3.09419 | 0.010689 | 0.018422 |
| hsa-miR-504-3p | -3.08889 | 0.002131 | 0.010115 |
| hsa-miR-643 | -3.08829 | 0.003244 | 0.011017 |
| hsa-miR-597-5p | -3.08748 | 0.002009 | 0.010115 |
| hsa-miR-532-3p | -3.0865 | 0.008232 | 0.015686 |
| hsa-miR-1248 | -3.08444 | 0.007967 | 0.015469 |
| hsa-miR-512-5p | -3.0826 | 0.004554 | 0.012360 |
| hsa-miR-212-3p | -3.08248 | 0.004314 | 0.012091 |
| hsa-miR-4521 | -3.08188 | 0.002205 | 0.010115 |
| hsa-miR-3613-3p | -3.07686 | 0.001822 | 0.010115 |
| hsa-miR-941 | -3.07015 | 0.000196 | 0.009233 |
| hsa-miR-345-3p | -3.06996 | 0.001070 | 0.009984 |
| hsa-miR-665 | -3.06002 | 0.001395 | 0.009984 |
| hsa-miR-758-3p+hsa-miR-411-3p | -3.05858 | 0.013977 | 0.021870 |
| hsa-miR-1323 | -3.0568 | 0.003475 | 0.011213 |
| hsa-miR-891a-5p | -3.05527 | 0.006132 | 0.013650 |
| hsa-miR-1269b | -3.05019 | 0.002648 | 0.010150 |
| hsa-miR-876-3p | -3.04605 | 0.005040 | 0.012745 |
| hsa-miR-548i | -3.04455 | 0.004292 | 0.012091 |
| hsa-miR-1206 | -3.04162 | 0.015355 | 0.023318 |
| hsa-miR-206 | -3.03896 | 0.005235 | 0.012837 |
| hsa-miR-579-5p | -3.03861 | 0.006237 | 0.013749 |
| hsa-miR-1203 | -3.02885 | 0.001419 | 0.009984 |
| hsa-miR-2110 | -3.02509 | 0.002629 | 0.010150 |
| hsa-miR-885-5p | -3.01969 | 0.008105 | 0.015623 |
| hsa-miR-103a-3p | -3.01938 | 0.002703 | 0.010150 |
| hsa-miR-450a-2-3p | -3.01645 | 0.001365 | 0.009984 |
| hsa-miR-3065-5p | -3.01578 | 0.008375 | 0.015838 |

| | | | |
|---|---|---|---|
| hsa-miR-520h | -3.01344 | 0.022590 | 0.031296 |
| hsa-miR-885-3p | -3.01218 | 0.024279 | 0.033290 |
| hsa-miR-10b-5p | -3.00318 | 0.001740 | 0.010115 |
| hsa-miR-1268b | -3.00239 | 0.001446 | 0.009984 |
| hsa-miR-1178-3p | -2.99787 | 0.000370 | 0.009488 |
| hsa-miR-548k | -2.9948 | 0.004379 | 0.012177 |
| hsa-mir-498 | -2.99466 | 0.002121 | 0.010115 |
| hsa-miR-584-3p | -2.99353 | 0.001476 | 0.009984 |
| hsa-miR-922 | -2.98964 | 0.007625 | 0.015328 |
| hsa-miR-210-3p | -2.98875 | 0.002158 | 0.010115 |
| hsa-miR-5010-3p | -2.98295 | 0.006406 | 0.014044 |
| hsa-miR-548g-3p | -2.98258 | 0.005047 | 0.012745 |
| hsa-miR-551b-3p | -2.97898 | 0.005233 | 0.012837 |
| hsa-miR-548a-5p | -2.97787 | 0.001210 | 0.009984 |
| hsa-miR-620 | -2.97721 | 0.008018 | 0.015493 |
| hsa-miR-373-3p | -2.97294 | 0.000995 | 0.009984 |
| hsa-miR-548m | -2.97288 | 0.004985 | 0.012745 |
| hsa-miR-490-3p | -2.96991 | 0.006792 | 0.014388 |
| hsa-miR-1281 | -2.96676 | 0.000644 | 0.009984 |
| hsa-miR-516b-5p | -2.9645 | 0.000676 | 0.009984 |
| hsa-miR-18b-5p | -2.96062 | 0.002857 | 0.010222 |
| hsa-miR-1245a | -2.96007 | 0.007704 | 0.015369 |
| hsa-miR-664a-3p | -2.95605 | 0.001018 | 0.009984 |
| hsa-miR-383-5p | -2.95525 | 0.000267 | 0.009488 |
| hsa-miR-574-3p | -2.95459 | 0.001268 | 0.009984 |
| hsa-miR-3164 | -2.95304 | 0.004084 | 0.011852 |
| hsa-miR-127-3p | -2.95261 | 0.004524 | 0.012321 |
| hsa-miR-324-3p | -2.94209 | 0.002619 | 0.010150 |
| hsa-miR-568 | -2.93982 | 0.001971 | 0.010115 |
| hsa-miR-146b-3p | -2.93775 | 0.004901 | 0.012745 |
| hsa-miR-449a | -2.93722 | 0.009201 | 0.016667 |
| hsa-mir-137 | -2.9353 | 0.005150 | 0.012837 |
| hsa-miR-892a | -2.93337 | 0.000749 | 0.009984 |
| hsa-miR-409-3p | -2.93022 | 0.005016 | 0.012745 |
| hsa-miR-624-3p | -2.92697 | 0.007221 | 0.014776 |
| hsa-miR-1976 | -2.92497 | 0.004265 | 0.012091 |
| hsa-miR-542-3p | -2.92491 | 0.001107 | 0.009984 |
| hsa-miR-449b-5p | -2.92199 | 0.007112 | 0.014708 |
| hsa-miR-548ai+hsa-miR-570-5p | -2.92154 | 0.007149 | 0.014708 |
| hsa-miR-449c-5p | -2.92059 | 0.025291 | 0.034149 |
| hsa-mir-375 | -2.91984 | 0.005487 | 0.013033 |

| | | | |
|---|---|---|---|
| hsa-miR-1291 | -2.91834 | 0.002848 | 0.010222 |
| hsa-miR-3605-5p | -2.91747 | 0.005521 | 0.013035 |
| hsa-miR-1255b-5p | -2.90198 | 0.001689 | 0.010115 |
| hsa-miR-378e | -2.90162 | 0.002672 | 0.010150 |
| hsa-miR-663a | -2.90032 | 0.011371 | 0.019255 |
| hsa-miR-297 | -2.90018 | 0.003659 | 0.011354 |
| hsa-miR-638 | -2.89688 | 0.011729 | 0.019664 |
| hsa-mir-520e | -2.89676 | 0.001712 | 0.010115 |
| hsa-miR-1469 | -2.89511 | 0.000951 | 0.009984 |
| hsa-mir-320a | -2.89427 | 0.016657 | 0.024799 |
| hsa-miR-520g-3p | -2.89395 | 0.005662 | 0.013097 |
| hsa-miR-302f | -2.89188 | 0.007097 | 0.014708 |
| hsa-miR-924 | -2.8911 | 0.000696 | 0.009984 |
| hsa-miR-151b | -2.8905 | 0.000475 | 0.009983 |
| hsa-miR-202-3p | -2.8888 | 0.005222 | 0.012837 |
| hsa-miR-299-5p | -2.88266 | 0.006671 | 0.014336 |
| hsa-miR-563 | -2.87882 | 0.008601 | 0.016036 |
| hsa-miR-802 | -2.87807 | 0.006454 | 0.014110 |
| hsa-miR-4792 | -2.87685 | 0.000445 | 0.009862 |
| hsa-miR-365a-3p+hsa-miR-365b-3p | -2.87679 | 0.009339 | 0.016822 |
| hsa-miR-506-5p | -2.87587 | 0.009259 | 0.016717 |
| hsa-miR-302a-3p | -2.86652 | 0.006683 | 0.014336 |
| hsa-miR-934 | -2.86445 | 0.006933 | 0.014521 |
| hsa-miR-572 | -2.86312 | 0.008521 | 0.015970 |
| hsa-miR-508-3p | -2.86088 | 0.003666 | 0.011354 |
| hsa-miR-1202 | -2.85183 | 0.001462 | 0.009984 |
| hsa-miR-4461 | -2.84901 | 0.002461 | 0.010150 |
| hsa-miR-99b-5p | -2.84757 | 0.005434 | 0.013033 |
| hsa-miR-1322 | -2.84646 | 0.000647 | 0.009984 |
| hsa-miR-935 | -2.84211 | 0.012372 | 0.020314 |
| hsa-miR-3151-5p | -2.84108 | 0.005638 | 0.013097 |
| hsa-miR-614 | -2.84 | 0.012929 | 0.020801 |
| hsa-miR-544a | -2.83998 | 0.018796 | 0.027123 |
| hsa-miR-371b-5p | -2.83968 | 0.007379 | 0.015040 |
| hsa-miR-664b-3p | -2.82929 | 0.003270 | 0.011041 |
| hsa-miR-188-3p | -2.82898 | 0.008159 | 0.015686 |
| hsa-miR-329-5p | -2.82635 | 0.020755 | 0.029263 |
| hsa-miR-573 | -2.82584 | 0.006930 | 0.014521 |
| hsa-miR-548h-5p | -2.82476 | 0.001653 | 0.010115 |
| hsa-miR-337-5p | -2.82108 | 0.001577 | 0.010115 |
| hsa-miR-944 | -2.82099 | 0.001414 | 0.009984 |

| | | | |
|---|---|---|---|
| hsa-miR-767-5p | -2.81934 | 0.008731 | 0.016202 |
| hsa-miR-199a-5p | -2.81401 | 0.007891 | 0.015451 |
| hsa-miR-219b-3p | -2.80778 | 0.036740 | 0.046612 |
| hsa-miR-4741 | -2.80539 | 0.004997 | 0.012745 |
| hsa-miR-3202 | -2.79878 | 0.032337 | 0.041689 |
| hsa-miR-219a-2-3p | -2.79704 | 0.013273 | 0.021141 |
| hsa-miR-1909-3p | -2.79683 | 0.004333 | 0.012091 |
| hsa-miR-376a-2-5p | -2.79405 | 0.010686 | 0.018422 |
| hsa-miR-378h | -2.78734 | 0.001661 | 0.010115 |
| hsa-mir-147b | -2.78674 | 0.005398 | 0.013033 |
| hsa-miR-553 | -2.78657 | 0.001336 | 0.009984 |
| hsa-miR-889-3p | -2.78558 | 0.001531 | 0.010115 |
| hsa-miR-509-5p | -2.78346 | 0.000331 | 0.009488 |
| hsa-miR-188-5p | -2.77943 | 0.012520 | 0.020432 |
| hsa-miR-5010-5p | -2.77651 | 0.021802 | 0.030416 |
| hsa-miR-483-3p | -2.77637 | 0.020271 | 0.028784 |
| hsa-miR-525-3p | -2.77483 | 0.015640 | 0.023548 |
| hsa-miR-154-5p | -2.77447 | 0.009192 | 0.016667 |
| hsa-miR-522-3p | -2.76767 | 0.008349 | 0.015825 |
| hsa-miR-376a-3p | -2.76609 | 0.000748 | 0.009984 |
| hsa-miR-1275 | -2.76318 | 0.001418 | 0.009984 |
| hsa-miR-548d-3p | -2.76309 | 0.005466 | 0.013033 |
| hsa-miR-1258 | -2.75866 | 0.002841 | 0.010222 |
| hsa-miR-28-3p | -2.75828 | 0.002596 | 0.010150 |
| hsa-miR-1244 | -2.75765 | 0.020030 | 0.028542 |
| hsa-miR-1257 | -2.75723 | 0.006037 | 0.013562 |
| hsa-miR-543 | -2.75522 | 0.010182 | 0.017779 |
| hsa-miR-936 | -2.75202 | 0.038136 | 0.048077 |
| hsa-miR-512-3p | -2.74845 | 0.002314 | 0.010150 |
| hsa-miR-4431 | -2.74567 | 0.002505 | 0.010150 |
| hsa-miR-554 | -2.74468 | 0.005285 | 0.012863 |
| hsa-miR-513c-3p | -2.73623 | 0.009178 | 0.016667 |
| hsa-miR-642a-3p | -2.72559 | 0.002208 | 0.010115 |
| hsa-miR-372-3p | -2.7234 | 0.018159 | 0.026540 |
| hsa-miR-548o-3p+hsa-miR-548ah-3p+hsa-miR-548av-3p | -2.7162 | 0.001476 | 0.009984 |
| hsa-miR-654-5p | -2.71618 | 0.007574 | 0.015301 |
| hsa-miR-218-5p | -2.71466 | 0.016648 | 0.024799 |
| hsa-miR-328-3p | -2.71404 | 0.007703 | 0.015369 |
| hsa-miR-4531 | -2.70667 | 0.015368 | 0.023318 |
| hsa-miR-532-5p | -2.70408 | 0.007138 | 0.014708 |
| hsa-miR-1271-3p | -2.70382 | 0.015586 | 0.023539 |

| | | | |
|---|---|---|---|
| hsa-miR-455-3p | -2.70034 | 0.000306 | 0.009488 |
| hsa-miR-329-3p | -2.69852 | 0.003700 | 0.011355 |
| hsa-miR-3161 | -2.69837 | 0.004214 | 0.012091 |
| hsa-miR-519d-3p | -2.69695 | 0.014113 | 0.021954 |
| hsa-miR-582-3p | -2.69426 | 0.001743 | 0.010115 |
| hsa-miR-139-5p | -2.69228 | 0.008525 | 0.015970 |
| hsa-miR-23b-3p | -2.69184 | 0.005339 | 0.012951 |
| hsa-miR-2278 | -2.68952 | 0.002514 | 0.010150 |
| hsa-miR-514b-3p | -2.68673 | 0.020198 | 0.028730 |
| hsa-miR-655-3p | -2.68663 | 0.001335 | 0.009984 |
| hsa-miR-548al | -2.68321 | 0.002544 | 0.010150 |
| hsa-miR-302d-3p | -2.67894 | 0.003796 | 0.011504 |
| hsa-miR-1279 | -2.6781 | 0.005169 | 0.012837 |
| hsa-miR-421 | -2.67543 | 0.000197 | 0.009233 |
| hsa-miR-890 | -2.67376 | 0.000142 | 0.008727 |
| hsa-miR-152-3p | -2.67215 | 0.000213 | 0.009421 |
| hsa-miR-296-3p | -2.66771 | 0.006797 | 0.014388 |
| hsa-miR-518f-3p | -2.66748 | 0.001387 | 0.009984 |
| hsa-miR-518d-3p | -2.66728 | 0.003485 | 0.011213 |
| hsa-miR-433-5p | -2.66695 | 0.026145 | 0.035007 |
| hsa-miR-376c-3p | -2.66549 | 0.006527 | 0.014232 |
| hsa-miR-1278 | -2.66349 | 0.003899 | 0.011610 |
| hsa-miR-4425 | -2.66156 | 0.004569 | 0.012360 |
| hsa-miR-128-3p | -2.66036 | 0.004516 | 0.012321 |
| hsa-miR-198 | -2.65995 | 0.017917 | 0.026331 |
| hsa-miR-761 | -2.65857 | 0.007169 | 0.014708 |
| hsa-miR-337-3p | -2.65776 | 0.003386 | 0.011041 |
| hsa-miR-190a-3p | -2.65546 | 0.009092 | 0.016667 |
| hsa-miR-589-5p | -2.64533 | 0.001246 | 0.009984 |
| hsa-miR-320d | -2.64504 | 0.001296 | 0.009984 |
| hsa-miR-326 | -2.64194 | 0.001951 | 0.010115 |
| hsa-miR-615-5p | -2.64182 | 0.007831 | 0.015430 |
| hsa-miR-765 | -2.64127 | 0.009178 | 0.016667 |
| hsa-miR-608 | -2.62839 | 0.026927 | 0.035728 |
| hsa-miR-551a | -2.62741 | 0.037015 | 0.046811 |
| hsa-mir-217 | -2.62475 | 0.018318 | 0.026627 |
| hsa-miR-127-5p | -2.62226 | 0.012295 | 0.020241 |
| hsa-miR-499b-3p | -2.62108 | 0.007991 | 0.015477 |
| hsa-miR-4707-3p | -2.61795 | 0.001836 | 0.010115 |
| hsa-miR-216b-5p | -2.61701 | 0.005596 | 0.013097 |
| hsa-miR-211-3p | -2.61674 | 0.026554 | 0.035376 |

| | | | |
|---|---|---|---|
| hsa-miR-511-5p | -2.61634 | 0.006207 | 0.013721 |
| hsa-miR-595 | -2.61409 | 0.013926 | 0.021833 |
| hsa-miR-516a-5p | -2.61215 | 0.004765 | 0.012611 |
| hsa-miR-4516 | -2.61071 | 0.009672 | 0.017189 |
| hsa-mir-378g | -2.60633 | 0.002742 | 0.010176 |
| hsa-miR-1302 | -2.60216 | 0.022890 | 0.031603 |
| hsa-miR-6511a-5p | -2.60169 | 0.001250 | 0.009984 |
| hsa-miR-371a-5p | -2.60068 | 0.007782 | 0.015426 |
| hsa-miR-151a-5p | -2.58881 | 0.008236 | 0.015686 |
| hsa-miR-1236-3p | -2.58621 | 0.012967 | 0.020821 |
| hsa-miR-3136-5p | -2.58487 | 0.003355 | 0.011041 |
| hsa-miR-598-3p | -2.58062 | 0.008462 | 0.015925 |
| hsa-miR-523-3p | -2.5773 | 0.006628 | 0.014335 |
| hsa-miR-374c-5p | -2.57437 | 0.010551 | 0.018303 |
| hsa-miR-1224-5p | -2.57219 | 0.002219 | 0.010115 |
| hsa-miR-3074-3p | -2.57184 | 0.024420 | 0.033425 |
| hsa-miR-1185-1-3p | -2.57154 | 0.018925 | 0.027211 |
| hsa-miR-613 | -2.56954 | 0.005619 | 0.013097 |
| hsa-miR-125a-5p | -2.56854 | 0.013601 | 0.021576 |
| hsa-miR-1226-3p | -2.56851 | 0.006681 | 0.014336 |
| hsa-miR-1287-5p | -2.56845 | 0.020698 | 0.029234 |
| hsa-miR-502-5p | -2.56742 | 0.008714 | 0.016202 |
| hsa-miR-3196 | -2.56582 | 0.009640 | 0.017189 |
| hsa-miR-183-5p | -2.5623 | 0.011356 | 0.019255 |
| hsa-miR-34b-3p | -2.56028 | 0.016131 | 0.024152 |
| hsa-miR-422a | -2.55615 | 0.003351 | 0.011041 |
| hsa-miR-519b-3p | -2.55485 | 0.007790 | 0.015426 |
| hsa-miR-526a+hsa-miR-518c-5p+hsa-miR-518d-5p | -2.55271 | 0.002067 | 0.010115 |
| hsa-miR-330-3p | -2.55128 | 0.003112 | 0.010763 |
| hsa-miR-410-3p | -2.54896 | 0.005506 | 0.013035 |
| hsa-miR-548e-3p | -2.54416 | 0.002698 | 0.010150 |
| hsa-miR-499b-5p | -2.54388 | 0.001164 | 0.009984 |
| hsa-miR-503-3p | -2.5422 | 0.007519 | 0.015267 |
| hsa-miR-671-3p | -2.54111 | 0.000520 | 0.009984 |
| hsa-miR-578 | -2.53821 | 0.006162 | 0.013659 |
| hsa-miR-548ak | -2.53743 | 0.001386 | 0.009984 |
| hsa-miR-138-5p | -2.53497 | 0.022675 | 0.031360 |
| hsa-miR-767-3p | -2.53492 | 0.010995 | 0.018795 |
| hsa-miR-331-5p | -2.5309 | 0.013642 | 0.021576 |
| hsa-miR-302e | -2.52725 | 0.030139 | 0.039299 |
| hsa-miR-1303 | -2.52694 | 0.000823 | 0.009984 |

| | | | |
|---|---|---|---|
| hsa-miR-891b | -2.52235 | 0.003621 | 0.011331 |
| hsa-miR-942-5p | -2.51628 | 0.004322 | 0.012091 |
| hsa-miR-518b | -2.51605 | 0.005705 | 0.013120 |
| hsa-miR-548j-3p | -2.51431 | 0.003506 | 0.011235 |
| hsa-miR-1268a | -2.51335 | 0.025230 | 0.034125 |
| hsa-miR-301b-3p | -2.51246 | 0.007546 | 0.015284 |
| hsa-miR-1287-3p | -2.5098 | 0.004807 | 0.012619 |
| hsa-miR-181b-2-3p | -2.50623 | 0.012804 | 0.020642 |
| hsa-miR-489-3p | -2.5051 | 0.012302 | 0.020241 |
| hsa-miR-182-5p | -2.50269 | 0.000693 | 0.009984 |
| hsa-let-7e-5p | -2.49452 | 0.002417 | 0.010150 |
| hsa-miR-501-3p | -2.48832 | 0.005553 | 0.013072 |
| hsa-miR-5196-3p+hsa-miR-6732-3p | -2.48656 | 0.033602 | 0.043110 |
| hsa-miR-587 | -2.48303 | 0.008934 | 0.016465 |
| hsa-miR-128-1-5p | -2.48224 | 0.008552 | 0.015982 |
| hsa-miR-208b-5p | -2.4782 | 0.004797 | 0.012619 |
| hsa-miR-1298-5p | -2.4682 | 0.004072 | 0.011852 |
| hsa-miR-378c | -2.4655 | 0.007597 | 0.015308 |
| hsa-miR-376c-5p | -2.46531 | 0.006014 | 0.013562 |
| hsa-miR-323b-5p | -2.46468 | 0.036899 | 0.046738 |
| hsa-miR-5196-5p | -2.46164 | 0.007155 | 0.014708 |
| hsa-miR-507 | -2.45883 | 0.025336 | 0.034153 |
| hsa-miR-509-3p | -2.44917 | 0.004443 | 0.012268 |
| hsa-miR-95-3p | -2.44428 | 0.005287 | 0.012863 |
| hsa-miR-450a-1-3p | -2.44357 | 0.006983 | 0.014588 |
| hsa-miR-494-3p | -2.44089 | 0.022343 | 0.031062 |
| hsa-miR-556-3p | -2.43897 | 0.004021 | 0.011780 |
| hsa-miR-758-5p | -2.43657 | 0.005792 | 0.013205 |
| hsa-miR-485-3p | -2.43493 | 0.003346 | 0.011041 |
| hsa-miR-196a-3p | -2.4335 | 0.011398 | 0.019255 |
| hsa-miR-494-5p | -2.42962 | 0.024759 | 0.033613 |
| hsa-miR-644a | -2.42857 | 0.008449 | 0.015925 |
| hsa-miR-128-2-5p | -2.42792 | 0.013088 | 0.020888 |
| hsa-miR-1306-3p | -2.42708 | 0.018682 | 0.027008 |
| hsa-miR-105-5p | -2.4215 | 0.006762 | 0.014388 |
| hsa-miR-299-3p | -2.41436 | 0.031233 | 0.040330 |
| hsa-miR-3140-5p | -2.40871 | 0.009210 | 0.016667 |
| hsa-miR-1289 | -2.40786 | 0.027904 | 0.036745 |
| hsa-miR-671-5p | -2.40562 | 0.028318 | 0.037167 |
| hsa-miR-519e-3p | -2.39514 | 0.024490 | 0.033464 |
| hsa-miR-1301-3p | -2.39323 | 0.014851 | 0.022872 |

| | | | |
|---|---|---|---|
| hsa-miR-561-3p | -2.38993 | 0.011191 | 0.019082 |
| hsa-miR-4421 | -2.38843 | 0.000896 | 0.009984 |
| hsa-miR-3144-3p | -2.38808 | 0.009762 | 0.017240 |
| hsa-miR-379-5p | -2.38707 | 0.012630 | 0.020526 |
| hsa-miR-874-5p | -2.38626 | 0.001367 | 0.009984 |
| hsa-miR-380-3p | -2.38575 | 0.009196 | 0.016667 |
| hsa-miR-610 | -2.38533 | 0.004930 | 0.012745 |
| hsa-miR-122-5p | -2.38381 | 0.009470 | 0.017020 |
| hsa-miR-1270 | -2.38353 | 0.017676 | 0.026025 |
| hsa-miR-640 | -2.38287 | 0.009871 | 0.017389 |
| hsa-miR-661 | -2.38173 | 0.018086 | 0.026482 |
| hsa-miR-320b | -2.37856 | 0.017051 | 0.025291 |
| hsa-miR-637 | -2.37813 | 0.014354 | 0.022199 |
| hsa-miR-1307-5p | -2.3769 | 0.004030 | 0.011780 |
| hsa-miR-3605-3p | -2.3756 | 0.031185 | 0.040330 |
| hsa-miR-606 | -2.36794 | 0.010017 | 0.017607 |
| hsa-miR-34c-3p | -2.36086 | 0.001209 | 0.009984 |
| hsa-miR-548ad-3p | -2.35739 | 0.019008 | 0.027281 |
| hsa-miR-590-3p | -2.35549 | 0.013442 | 0.021368 |
| hsa-miR-873-5p | -2.35091 | 0.014876 | 0.022872 |
| hsa-miR-3916 | -2.35075 | 0.012125 | 0.020116 |
| hsa-miR-2116-5p | -2.3507 | 0.010662 | 0.018422 |
| hsa-miR-627-5p | -2.34541 | 0.013720 | 0.021595 |
| hsa-miR-612 | -2.34489 | 0.005424 | 0.013033 |
| hsa-miR-516a-3p+hsa-miR-516b-3p | -2.34278 | 0.002123 | 0.010115 |
| hsa-miR-1827 | -2.34073 | 0.008771 | 0.016239 |
| hsa-miR-520a-5p | -2.33694 | 0.026296 | 0.035149 |
| hsa-miR-651-5p | -2.33542 | 0.002955 | 0.010443 |
| hsa-miR-3131 | -2.32982 | 0.019345 | 0.027715 |
| hsa-miR-362-5p | -2.32973 | 0.007925 | 0.015451 |
| hsa-miR-539-5p | -2.32879 | 0.015433 | 0.023369 |
| hsa-miR-181a-3p | -2.3278 | 0.021703 | 0.030331 |
| hsa-miR-515-3p | -2.3264 | 0.017618 | 0.025988 |
| hsa-mir-520b | -2.32481 | 0.011484 | 0.019293 |
| hsa-miR-320c | -2.32329 | 0.014766 | 0.022792 |
| hsa-miR-192-5p | -2.32081 | 0.001154 | 0.009984 |
| hsa-miR-3180 | -2.32062 | 0.003115 | 0.010763 |
| hsa-miR-1245b-5p | -2.31892 | 0.009516 | 0.017065 |
| hsa-miR-1285-3p | -2.31747 | 0.015275 | 0.023306 |
| hsa-miR-4435 | -2.30675 | 0.030492 | 0.039630 |
| hsa-miR-564 | -2.29948 | 0.039250 | 0.049093 |

| | | | |
|---|---|---|---|
| hsa-miR-3180-5p | -2.29937 | 0.002605 | 0.010150 |
| hsa-miR-548d-5p | -2.29776 | 0.000341 | 0.009488 |
| hsa-miR-1204 | -2.2956 | 0.021102 | 0.029699 |
| hsa-miR-96-5p | -2.29243 | 0.017211 | 0.025481 |
| hsa-miR-382-5p | -2.29064 | 0.007828 | 0.015430 |
| hsa-miR-1910-3p | -2.28875 | 0.013019 | 0.020861 |
| hsa-miR-3180-3p | -2.28704 | 0.003321 | 0.011041 |
| hsa-miR-376b-3p | -2.2868 | 0.014319 | 0.022187 |
| hsa-miR-1299 | -2.28366 | 0.013695 | 0.021595 |
| hsa-miR-365b-5p | -2.28347 | 0.009655 | 0.017189 |
| hsa-miR-744-5p | -2.27832 | 0.019693 | 0.028113 |
| hsa-miR-411-5p | -2.27672 | 0.030626 | 0.039688 |
| hsa-miR-381-3p | -2.27385 | 0.037705 | 0.047608 |
| hsa-miR-298 | -2.2693 | 0.036329 | 0.046178 |
| hsa-miR-382-3p | -2.26866 | 0.005790 | 0.013205 |
| hsa-miR-708-5p | -2.26649 | 0.003887 | 0.011610 |
| hsa-miR-204-5p | -2.26446 | 0.007388 | 0.015040 |
| hsa-miR-542-5p | -2.24949 | 0.006578 | 0.014303 |
| hsa-miR-548l | -2.24502 | 0.007938 | 0.015451 |
| hsa-miR-193a-5p+hsa-miR-193b-5p | -2.24138 | 0.005739 | 0.013160 |
| hsa-miR-31-5p | -2.23234 | 0.027197 | 0.035933 |
| hsa-miR-367-3p | -2.23146 | 0.013048 | 0.020867 |
| hsa-miR-539-3p | -2.23032 | 0.004277 | 0.012091 |
| hsa-miR-195-5p | -2.22385 | 0.007687 | 0.015369 |
| hsa-miR-221-3p | -2.22182 | 0.000472 | 0.009983 |
| hsa-miR-302c-3p | -2.21942 | 0.023251 | 0.031990 |
| hsa-miR-1273c | -2.21934 | 0.038538 | 0.048430 |
| hsa-miR-362-3p | -2.21134 | 0.002800 | 0.010222 |
| hsa-miR-3192-5p | -2.2107 | 0.003994 | 0.011761 |
| hsa-miR-504-5p | -2.20302 | 0.024647 | 0.033564 |
| hsa-miR-629-5p | -2.20059 | 0.012183 | 0.020170 |
| hsa-miR-892b | -2.20024 | 0.016020 | 0.024030 |
| hsa-miR-1307-3p | -2.19987 | 0.004983 | 0.012745 |
| hsa-miR-1264 | -2.19597 | 0.014299 | 0.022187 |
| hsa-miR-370-5p | -2.1914 | 0.001363 | 0.009984 |
| hsa-miR-491-3p | -2.19123 | 0.018857 | 0.027162 |
| hsa-miR-675-5p | -2.18838 | 0.017951 | 0.026333 |
| hsa-miR-591 | -2.18665 | 0.026445 | 0.035290 |
| hsa-miR-4458 | -2.18407 | 0.023614 | 0.032433 |
| hsa-miR-1-5p | -2.17698 | 0.015019 | 0.023023 |
| hsa-miR-3918 | -2.17142 | 0.021583 | 0.030269 |

| | | | |
|---|---|---|---|
| hsa-miR-3615 | -2.16219 | 0.005241 | 0.012837 |
| hsa-miR-1306-5p | -2.16217 | 0.003176 | 0.010832 |
| hsa-miR-2053 | -2.16055 | 0.011459 | 0.019292 |
| hsa-miR-101-3p | -2.14936 | 0.026613 | 0.035396 |
| hsa-miR-330-5p | -2.13242 | 0.002429 | 0.010150 |
| hsa-miR-641 | -2.13002 | 0.005680 | 0.013100 |
| hsa-miR-92b-3p | -2.1297 | 0.024533 | 0.033466 |
| hsa-miR-125a-3p | -2.12874 | 0.025462 | 0.034264 |
| hsa-miR-1183 | -2.12454 | 0.021163 | 0.029732 |
| hsa-miR-147a | -2.12248 | 0.024768 | 0.033613 |
| hsa-miR-203a-3p | -2.11647 | 0.029602 | 0.038726 |
| hsa-miR-384 | -2.11423 | 0.016643 | 0.024799 |
| hsa-miR-1304-3p | -2.11065 | 0.026146 | 0.035007 |
| hsa-miR-617 | -2.10883 | 0.038818 | 0.048629 |
| hsa-miR-1293 | -2.1063 | 0.005237 | 0.012837 |
| hsa-miR-1908-5p | -2.09696 | 0.012712 | 0.020576 |
| hsa-miR-485-5p | -2.09334 | 0.039425 | 0.049235 |
| hsa-miR-520d-5p+hsa-miR-527+hsa-miR-518a-5p | -2.0862 | 0.002619 | 0.010150 |
| hsa-miR-377-3p | -2.08609 | 0.025013 | 0.033888 |
| hsa-miR-660-5p | -2.07701 | 0.002489 | 0.010150 |
| hsa-miR-4524a-5p | -2.04043 | 0.012664 | 0.020540 |
| hsa-miR-378b | -2.02331 | 0.010917 | 0.018735 |
| hsa-miR-4286 | -2.02301 | 0.033099 | 0.042533 |
| hsa-miR-132-3p | -2.02051 | 0.006141 | 0.013650 |
| hsa-miR-571 | -2.01413 | 0.033780 | 0.043268 |
| hsa-miR-631 | -2.01406 | 0.018459 | 0.026753 |
| hsa-miR-221-5p | -1.98034 | 0.039764 | 0.049581 |
| hsa-miR-339-5p | -1.97199 | 0.014108 | 0.021954 |
| hsa-miR-605-5p | -1.96587 | 0.036330 | 0.046178 |
| hsa-miR-186-5p | -1.95828 | 0.012298 | 0.020241 |
| hsa-miR-197-3p | -1.94121 | 0.032394 | 0.041695 |
| hsa-miR-345-5p | -1.92914 | 0.021700 | 0.030331 |
| hsa-miR-200c-3p | -1.92885 | 0.029106 | 0.038139 |
| hsa-miR-3928-3p | -1.9192 | 0.027602 | 0.036407 |
| hsa-miR-6511a-3p | -1.87652 | 0.036340 | 0.046178 |
| hsa-miR-185-5p | -1.87369 | 0.022404 | 0.031093 |
| hsa-miR-496 | -1.86714 | 0.038720 | 0.048583 |
| hsa-miR-619-3p | -1.86042 | 0.030443 | 0.039630 |
| hsa-mir-190b | -1.83281 | 0.021893 | 0.030490 |
| hsa-miR-596 | -1.83164 | 0.023079 | 0.031809 |
| hsa-miR-576-5p | -1.77425 | 0.034996 | 0.044754 |

| | | | |
|---|---|---|---|
| hsa-miR-323b-3p | -1.76895 | 0.029956 | 0.039124 |
| hsa-miR-193b-3p | -1.7548 | 0.036279 | 0.046178 |
| hsa-miR-342-5p | -1.74782 | 0.010917 | 0.018735 |
| hsa-miR-503-5p | -1.66492 | 0.038407 | 0.048342 |
| hsa-miR-6724-5p | -1.57389 | 0.030636 | 0.039688 |
| hsa-miR-1180-3p | -1.55797 | 0.015370 | 0.023318 |
| hsa-miR-374b-5p | -1.54354 | 0.015892 | 0.023883 |
| hsa-miR-1296-5p | -2.212 | 0.020390 | 0.028900 |
| hsa-miR-3613-5p | -2.649 | 0.002664 | 0.010150 |
| hsa-miR-490-5p | -2.738 | 0.017038 | 0.025291 |
| hsa-miR-887-3p | -2.774 | 0.005244 | 0.012837 |
| hsa-miR-203a-5p | -2.923 | 0.005440 | 0.013033 |

Supplementary table 3 qPCR analysis for hsa-miR-223-3p, hsa-miR-150-5p, hsa-miR-100-5p and hsa-miR-494-3p against the endogenous control RNU6B

| | hsa-miR-100-5p | hsa-miR-150-5p | hsa-miR-223-3p | hsa-miR-494-3p | |
|---|---|---|---|---|---|
| **TC-39** | 0,06 | 79,66 | 201,24 | 3,46 | BuAL |
| **TC-41** | 0,35 | 387,13 | 759,15 | 7,53 | BuAL |
| **TC-43** | 0,45 | 987,84 | 4958,54 | 6,95 | BuAL |
| **TC-44** | 0,15 | 126,03 | 629,39 | 0,76 | BuAL |
| **TC-45** | 0,37 | 289,85 | 3465,21 | 4,00 | BuAL |
| **TC-46** | 3,70 | 1279,73 | 1195,26 | 7,57 | BuAL |
| **TC-49** | 0,17 | 118,19 | 1738,50 | 8,14 | BuAL |
| **TC-38** | 21,05 | 160,01 | 350,59 | 6,94 | BuLY |
| **TC-40** | 11,87 | 820,80 | 468,93 | 29,71 | BuLY |
| **TC-51** | 0,09 | 39,43 | 41,88 | 8,40 | BuLY |
| **TC-52** | 1,04 | 10,06 | 6,31 | 20,62 | BuLY |
| **TC-53** | 0,58 | 111,68 | 19,44 | 4,86 | BuLY |
| **TC-54** | 0,11 | 340,46 | 5224,48 | 2,54 | BuLY |
| **TC-76** | 2,11 | 4511,83 | 1567,74 | 4,2 | Norm LN |
| **TC-77** | 2,77 | 2518,77 | 236,74 | 3,78 | Tonsil |
| **TC-78** | 1,9 | 1743,18 | 241,63 | 5,06 | Tonsil |
| **TC-80** | 2,17 | 3650,95 | 1059,84 | 7,02 | NCLT Mix |
| **TC-81** | 2,15 | 3049,72 | 511,54 | 4,92 | NCLT Mix |
| **DG-75** | 4,75 | 10,27 | 0,73 | 2,95 | BL cell line |
| **Raji** | 44,52 | 0,03 | 10,08 | 6,57 | BL cell line |
